# 3D Printers in Hospitals: Bacterial Contamination of Common and Antimicrobial 3D-Printed Material

**DOI:** 10.1101/2024.03.30.587440

**Authors:** Katelin C. Jackson, Erin Clancey, Douglas R. Call, Eric Lofgren

## Abstract

COVID-19 has presented hospitals with unique challenges. A SHEA Research Network survey showed that 40% reported “limited” or worse levels of personal protective equipment (PPE), and 13% were self-producing PPE to address those deficits, including 3D-printed items. However, we do not know how efficiently, if at all, 3D-printed materials can be disinfected. Additionally, two filaments, PLACTIVE and BIOGUARD, claim to be antimicrobial; they use copper nanocomposites and silver ions to reduce bacterial populations. We assess how PLACTIVE and BIOGUARD may be contaminated and how well they reduce contamination, and how readily Polylactic Acid (PLA), a standard 3D-printed material, may be disinfected.

3D-printed materials, including PLACTIVE and BIOGUARD, are readily contaminated with bacteria that are common in hospitals and can sustain that contamination. Our findings reveal that the levels of contamination on PLACTIVE and BIOGUARD can vary under specific conditions such as layer height or bacterial contact time, sometimes surpassing or falling short of PLA. However, disinfected disks had lower overall CFU averages than those that were not, but the level of disinfection was variable, and bacterial populations recovered hours after disinfection application. Proper disinfection and using appropriate 3D-printed materials are essential to limit bacterial contamination. 3D printers and their products can be invaluable for hospitals, especially when supplies are low, and healthcare worker safety is paramount. Environmental services should be made aware of the presence of antimicrobial 3D-printed materials, and patients should be discouraged from printing their own items for use in hospital environments.

**IMPORTANCE:** The COVID-19 pandemic has intensified the demand for personal protective equipment (PPE) in hospitals, prompting the utilization of 3D-printed materials to address shortages. Given the role of environmental contamination in healthcare-associated infections, understanding the potential for bacterial colonization on these materials is crucial. Our findings highlight the importance of proper disinfection practices and material selection in mitigating bacterial contamination, enhancing infection prevention strategies in hospitals, and ensuring the safety of healthcare workers and patients.

## INTRODUCTION

An estimated 20% to 40 % of healthcare-associated infections (HAIs) have been attributed to cross infections from the hands of healthcare workers (HCWs), who have become contaminated either directly by patient contact or indirectly via touching of contaminated surfaces [1], [2]. It has been strongly suggested that environmental contamination plays a vital role in pathogen transmissions such as methicillin-resistant *Staphylococcus aureus* (MRSA) and vancomycin-resistant Enterococcus spp. Recent evidence suggests this environmental transmission is linked to other nosocomial infections such as Norovirus, *Clostridium difficile*, and Acinetobacter spp. [1], [3]. Many gram-negative bacteria can survive for long periods on environmental surfaces and can be a continual transmission source. A variable degree of pathogen transfer can come from single-hand contact with a contaminated surface, and contaminated hands can transfer bacteria and viruses to other surfaces and subjects [4].

Antimicrobial-resistant bacteria have prompted the exploration of alternative solutions, such as utilizing silver, copper, or copper alloys either as coatings on surfaces like stainless steel or as standalone materials, to mitigate bacterial pathogens in hospital settings [5], [6], [7]. Both silver and copper have been employed for centuries as antimicrobial agents, owing to their effectiveness in neutralizing bacteria through various mechanisms [8], [9]. Silver, in particular, exhibits high toxicity towards bacteria, employing multiple mechanisms for bacterial inactivation [7], [8], [10]. Silver ions have been shown to inhibit the respiratory chain in *Escherichia coli* [11], inhibit enzymes, interfere with electron transport, and bind to DNA [12], [13]. Silver nanoparticles are believed to cause cell membrane leakage and penetrate the cell, leading to inhibition of several cell processes, i.e., transcription, translation, and protein synthesis, and impairment of essential cellular function [14], [15]. Copper and its alloys have been registered by the US Environmental Protection Agency (EPA) as the first solid antimicrobial material [16] [17]. While copper ions are essential for bacteria, uncontrolled levels of free ions can lead to toxic cellular effects [18]. Copper’s mechanism of action is thought to be cell membrane degradation by copper ions that are released from the copper surfaces, allowing the copper ions to penetrate the cell, leading to oxidative stress and DNA damage [9], [19].

Since the beginning of the COVID-19 pandemic, anecdotal reports have indicated hospitals have faced supply shortages, i.e., hand hygiene/disinfectant, N95 respirators, ventilator parts, etc., leaving healthcare workers to reuse personal protection equipment (PPE). A survey of hospitals showed that 40% reported “limited” or worse levels of PPE, and 13% were self-producing PPE to address those deficits [20]. This shortage has introduced a novel type of surface into the healthcare environment: 3D-printed material. To address the shortage, hospitals have accepted 3D-printed PPE and other necessary supplies from sources, i.e., tech companies, individuals with 3D printers, etc., willing to make and donate these supplies. The influx of outside support had led to a “COVID-19 Supply Chain response curated by the NIH/NIAID in collaboration with the US FDA, the Veterans Healthcare Administrations, and America Makes to support the manufacturing of PPE and other medical equipment” [21]. This collaboration can verify these designs’ quality, safety, and efficacy. We do not know how efficiently, if at all, how these 3D-printed materials can withstand hospital infection prevention practices.

This study aims to determine if 3D-printed material can be contaminated with hospital-associated bacteria, as well as how effectively if at all, these same materials can be disinfected. We assess how readily the most common material used in small-scale 3D printing, Polylactic Acid (PLA), may be both contaminated and disinfected. Additionally, two PLA-based filaments, PLACTIVE and BIOGUARD, with antimicrobial copper nanocomposites and silver additives, respectively, are evaluated for their ability to reduce bacterial contamination.

## METHODS

### Materials

We used 3D-printed 30mm diameter discs, 5mm in height, with a small additional 3mm bar on top of the disk to facilitate gripping, as our experimentally contaminated surface (model file available at https://github.com/epimodels/3dprinted_PPE). These disks were printed using a Prusa Research Mk3S model 3D printer in an environment intended to mimic a home or maker space environment (i.e., not under sterile conditions). We used three different polylactic acid filaments: unmodified Natureworks Ingeo 4043D PLA (Protoplant, Vancouver, WA), PLACTIVE (Copper3D, Santiago, Chile), and BIOGUARD (3DXTECH, Grand Rapids, MI). PLA is a commonly used material, and while PLACTIVE and BIOGUARD are less common, they claim to be antimicrobial due to the added copper nanocomposites and silver ions, respectively. In our experimentation, we considered PLA as a reference material. All disks were printed using two layer heights, 0.2mm and 0.3mm, to see if variation in thickness affects the amount of bacterial growth. These heights were chosen as common heights for detailed vs. fast printing, respectively. Before experimentation, all 3D-printed disks were sterilized. Disks were placed in a container of 95% EtOH and shaken for 10 minutes. Afterward, they underwent UV sterilization for 1 hour per side.

### Bacterial Growth and Viability

We performed a time series experiment to determine bacterial growth and viability on each of our filament materials. Specifically, we grew methicillin-resistant *Staphylococcus aureus* (ATCC BAA 1747; ATCC 43300), *Staphylococcus aureus* (ATCC 25923; ATCC 29213), *Escherichia coli* (AR-0450, AR-0373), and *Klebsiella pneumoniae* (AR-0087, AR-0115) on PLA, PLACTIVE, and BIOGUARD. Bacteria were diluted to a concentration of 0.06 per mL (10^7^). We then pipetted 25ul (between 10^5^ and 10^6^) onto each disk, spread the droplet with an insulating loop, and then dried each disk under the hood for short (3 hours) and long (24 hours) contact times. Afterward, the disks were placed in 6-well plates with 1.5 ml tryptic soy broth (TSB) media and put into the incubator at 37° C for a time course of 0-, 4-, 6-, 8-, 12-, and 16-hours. At each time point, 1:10 serial dilutions in TSB were performed to attain colony-forming unit (CFU) counts for each strain per disk. We then pipetted 5ul of diluted bacteria onto tryptic soy agar (TSA) plates in triplicate for replication. We ran each experiment in technical duplicates. Plates were incubated at 37°C overnight. We counted the number of CFUs occurring on each plate. The CFU counts from the reference group (PLA) and testing group (PLACTIVE and BIOGUARD) were analyzed to see how well the antimicrobial material decreased bacterial load.

### Disinfection

To determine if 3D-printed material can be disinfected, we used 70% EtOH on PLA only. After the disks were dried, we sprayed 70% EtOH onto each disk for full coverage, then left them to dry under the hood for 30 seconds with wiping, 2.5 mins, 5 mins, and 10 mins. Disks were placed in 6-well plates with 1.5 ml TSB media and placed into the incubator at 37° C for a time course of 0-, 4-, 6-, 8-, and 16-hours. For each time point, 1:10 serial dilutions in TSB were performed to attain the disinfectant PLA CFU counts for each strain per disk. We pipetted 5ul of diluted bacteria onto TSA plates in triplicate for replication, and the experiment was run in technical duplicates. Plates were placed in the 37°C incubator overnight. The CFU counts of PLA with and without disinfectant were analyzed to see how well 70% EtOH decreased bacterial load.

### Statistical Analysis

We conducted a multifactorial analysis of variance (ANOVA) to assess the factors affecting CFU counts. Then, we used nonlinear least squares (NLS) to estimate the growth curves of the different bacterial strains of each material across all time points. Bacterial strains were combined under the corresponding bacterial species for the analysis. The multifactorial ANOVA enabled us to assess the significance of CFU counts across various experimental conditions. We used material type, bacterial strains, layer height, bacterial contact time, disinfectant contact times, and the interactions between these variables to predict CFU counts. Post-hoc comparisons were subsequently conducted to investigate pairwise differences.

Next, we employed NLS to fit a sigmoid function to describe the bacterial growth curve. The sigmoid function, also known as the logistic growth model, is given by:

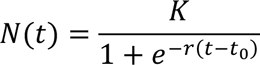

where *N*(*t*) represents the population size (in this case, the CFU counts) at time *t*, *K* is the carrying capacity, *r* is the intrinsic growth rate, *t* is the time variable, and *t*_0_ is the time at which the population size is at its initial value *N*_0_.

We used NLS to fit this sigmoid function to our experimental data, estimating the parameters *r*, *K*, and *N*_0_. Starting parameter values were *r* = 0.5, *K* = 500000000, and *N*_0_ = 40000. Although our response variable, CFU, is given in counts, nonlinear regression models assume data is continuous but are nonetheless appropriate for count data [22], [23]. These parameter estimates provided insights into the growth characteristics of the bacterial strains under different experimental conditions. Statistical analysis was done in the R Statistical Software language using the minpack.lm library [24], [25]. Data and analytical code is available at https://github.com/epimodels/3dprinted_PPE.

Following model fitting, we examined various experimental conditions, including short (3-hour) and long (24-hour) bacterial contact times, different layer heights, PLA with and without disinfectant, and different material types by calculating the differences in growth rate and carrying capacity. Subsequently, we utilized standard error to compute 95% confidence intervals (CI) for the comparison of each model.

### Scanning Electron Microscopy (SEM) Materials and Methods

Samples were brought to the Franceschi Microscopy and Imaging Center in 2% paraformaldehyde, 2% glutaraldehyde, 0.1M Phosphate buffer after an overnight incubation overnight at 4° C. Samples were rinsed twice with 0.1M Phosphate buffer then post-fixed overnight in 1% osmium tetroxide at 4° C. After water rinses, they were dehydrated with an EtOH series (30% -100%). The final drying was with hexamethyldisilizane (HMDS). Samples were mounted on aluminum stubs and gold coated. Due to damage from the electron beam, samples were removed from the stubs, the underside was gold coated, then samples were remounted and gold coated again. Samples were then placed in the SEM, and double sticky carbon tape was placed across the surface in two places. Then, it was attached to the SEM stage. Images are shown in Figure 1.

**Figure 1.**
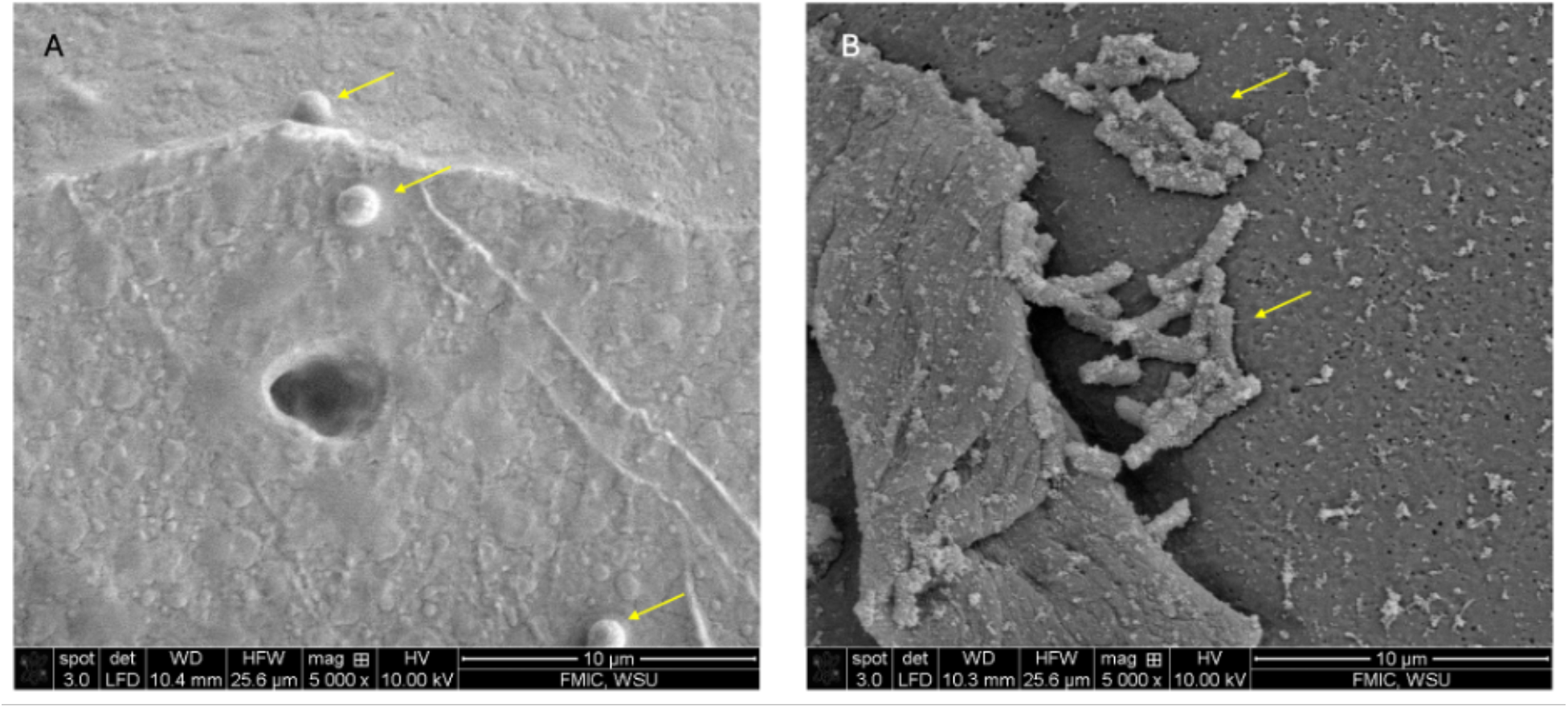
Scanning electron micrograph of (A) MRSA on PLA 0.2mm and (B) Klebsiella pneumonia on PLA 0.2mm, located within the groves of the PLA disk. Samples were fixed with 2% paraformaldehyde, 2% glutaraldehyde, 0.1M Phosphate buffer overnight at 4° C, rinsed twice with 0.1M Phosphate buffer, and then post-fixed overnight in 1% osmium tetroxide at 4° C. After water rinses, they were dehydrated with an EtOH series (30% -100%) followed by final drying with hexamethyldisilizane (HMDS). Samples were mounted on aluminum stubs, gold-coated twice to mitigate electron beam damage, and imaged with a FEI Quanta 200F SEM.

## RESULTS

### Multifactorial Analysis of Bacterial Contamination

To address the study’s central question, which was whether bacterial contamination occurs on polylactic acid (PLA) and if this material can be effectively disinfected. A multifactorial analysis of variance (ANOVA) was conducted to assess the effects of material type, bacteria, layer height, bacterial contact time, and disinfectant contact times on CFU. The ANOVA revealed significant effects of material type (F(2, 3448) = 39.6525, p < 0.001) and Bacteria (F(3, 3448) = 14.8295, p < 0.001) on Colony Forming Units (CFU).

Post-hoc comparisons revealed significant differences in CFU between PLA and PLACTIVE (t-values = -2.703, p = 0.0069), as well as between PLA and BIOGUARD (t-value = 5.997, p < 0.001), indicating variations in bacterial growth across different material types. Furthermore, the coefficients for methicillin-resistant *Staphylococcus aureus* and *Staphylococcus aureus* suggest that the mean CFU significantly differs from the reference species, *E. coli*. They had statistically significant negative effects on CFU compared to *E. coli* (t-values = -11.726, -12.436 respectively, p < 0.001. Meanwhile, *Klebsiella pneumoniae* was different but nonsignificant (t-value = -0.584, p = 0.5593).

Investigating the effectiveness of disinfection on polylactic acid (PLA), the analysis focused explicitly on PLA with a layer height of 0.2 mm and a 3-hour bacterial contact time. Specifically, in the absence of disinfectant, we found a significantly higher mean CFU than when using 70% EtOH (estimate = 725976000, t-value = 13.071, p < 0.001). Additionally, it is noteworthy that neither layer height, bacterial contact time, nor disinfectant contact times showed significant effects on CFU (layer height: F(1, 3448) = 0.6484, p = 0.4207; bacterial contact time: F(1, 3448) = 0.0094, p = 0.9227; disinfectant contact time: F(3, 1192) = 1.4284, p = 0.2328), suggesting that variations in these factors did not significantly impact bacterial contamination levels in the context of this study.

However, further analysis was conducted using non-linear models to explore these possible variations. This additional investigation aimed to uncover differences in growth rate (r) and carrying capacity (K) between Material Types, assess the impact of bacterial contact time on CFU, and investigate the influence of layer height on bacterial contamination. These analyses were pursued to deepen our understanding of the factors contributing to bacterial contamination in the context of the study.

### Material Type Impact on Growth Rate and Carrying Capacity

When investigating the difference in growth rate (r) and carrying capacity (k) between material types, we found that for *Klebsiella pneumonia*, there was a decrease in carrying capacity between PLA and BIOGUARD for the long bacterial contact time (24hr) at 0.2mm (-6.64e9 [-1.15e10, -1.74e9]) and 0.3mm (-1.033e9 [-1.76e9, -3.05e8]), with PLA having lower CFU counts than BIOGUARD. This also occurred for PLA and BIOGAURD for the short bacterial contact time (3 hours) at 0.3mm (-2.24e9 [-3.13e9, - 1.36e9]). On the other hand, there was an increase in carrying capacity between PLA and PLACTIVE for the short bacterial contact time (3 hours) at 0.3mm (5.82e8 [8.85e7, 1.08e9]), with PLA having higher CFU counts than PLACTIVE.

We found a higher growth rate of *Escherichia coli* for PLA when compared to PLACTIVE for the long bacterial contact time (24 hours) at 0.2mm (1.08 [0.427, 1.74]). Additionally, there was a decrease in carrying capacity between PLA and BIOGUARD for 24-hour contact time at 0.2mm (-1.55e9 [-2.40e9, -6.86e8]) and for 3-hour contact time at 0.3mm (-1.86e9 [-2.45e9, -1.28e9]) and between PLA and PLACTIVE for 3 hour contact time at 0.3mm (-5.69e8 [-1.05e9, -8.58e7]), with PLA having lower CFU counts.

Methicillin-resistant *Staphylococcus aureus* had an increased carrying capacity for PLA when compared to PLACTIVE for both long (5.25e8 [1.54e8, 8.97e8]) and short (1.58e9 [1.08e9, 2.09e9]) bacterial contact times at 0.2mm and 0.3mm, respectively. However, PLA had a decrease in carrying capacity when compared to BIOGUARD for the long 24-hour bacterial contact time at 0.3mm (-2.65e9 [-4.24e9, -1.05e9]) but showed an increase in carrying capacity for the short 3-hour bacterial contact time at 0.3mm (1.12e9 [3.7e8, 1.9e9]). (Figure 2 & Figure 3)

**Figure 2:**
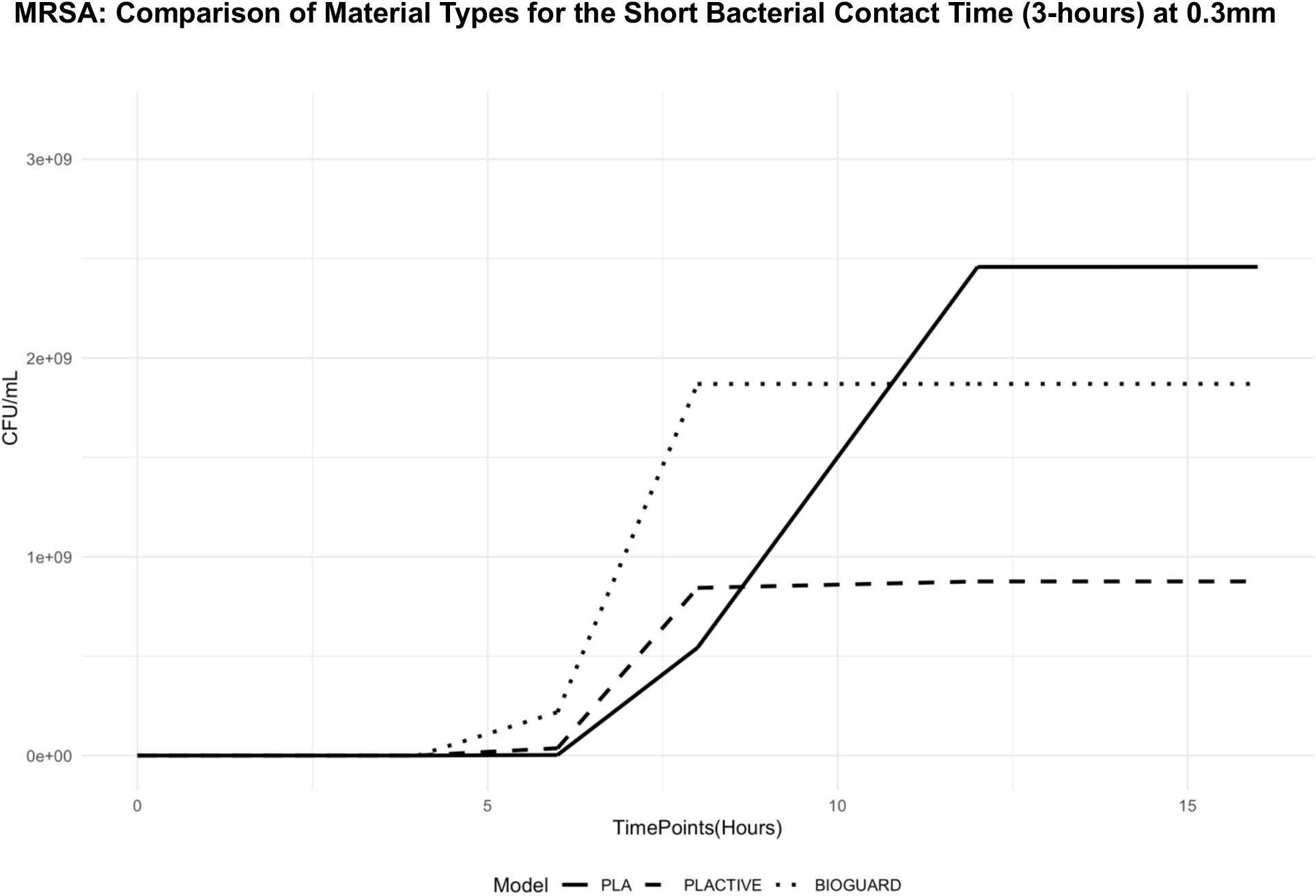
Growth curve of methicillin-resistant *Staphylococcus aureus,* comparing CFU/ml of the three material types, PLA, PLACTIVE, and BIOGUARD, for the short bacterial contact time at 0.3mm.

**Figure 3:**
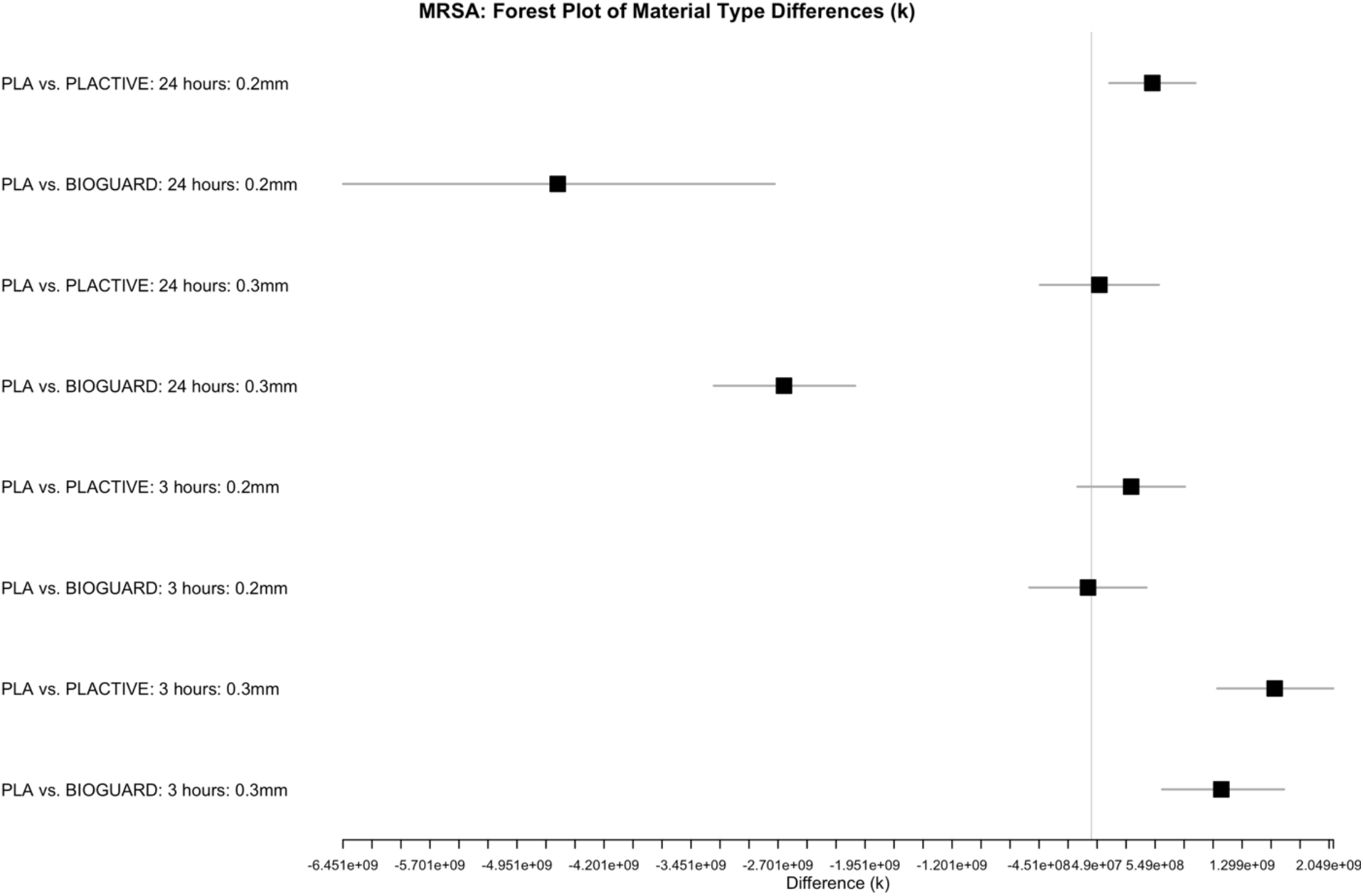
Forest plot illustrating the differences in k values between material types for methicillin-resistant *Staphylococcus aureus,* categorized by contact time and layer height. The plot displays the mean difference (k) along with scaled confidence intervals (CIs), where CIs exceeding a threshold value are scaled by a specified factor.

Lastly, there was a decrease in carrying capacity for *Staphylococcus aureus* between PLA and BIOGUARD for the 24-hour bacterial contact time at both 0.2mm (-1.36e9 [-2.0e9, -7.17e8]) and 0.3mm (-1.14e9 [-1.61e9, -6.71e8]). However, there was an increase in carrying capacity for PLA and PLACTIVE for the 3-hour bacterial contact time at 0.3mm (6.69e8 [1.87e8, 1.15e9]), with CFU counts being higher for PLA.

Our findings reveal variability in bacterial CFU levels between materials, with PLA exhibiting either higher or lower bacterial CFU than the alternative antimicrobial materials PLACTIVE and BIOGUARD, depending on various factors. Critically, there is no apparent consistent trend between standard PLA and materials containing antimicrobial additives (Supplement Figures 1-7).

### Contact Time Impact on Growth Rate and Carrying Capacity

Our models show a notable difference between the long (24 hours) and short (3 hours) bacterial contact times, with the 24-hour contact time consistently exhibiting higher CFU/mL compared to the shorter 3-hour contact time for all bacteria tested except for one case of MRSA where the carrying capacity decreased for the long bacterial contact time when using PLA 0.3mm (-1.18e9 [-1.72e9, -6.49e8]). With *Klebsiella pneumonia* using PLA 0.3mm, a significant increase in carrying capacity was observed, resulting in higher CFU counts (1.59e9 [1.03e9, 2.15e9]) for the longer bacterial contact time. Similar trends were observed with PLACTIVE 0.3mm (2.17e9 [1.42e9, 2.91e9]). Furthermore, BIOGUARD 0.2mm demonstrated an increased carrying capacity for the longer 24-hour contact time (6.41e9 [1.51e9, 1.13e10]) compared to the shorter 3-hour contact time. (Figure 4)

**Figure 4:**
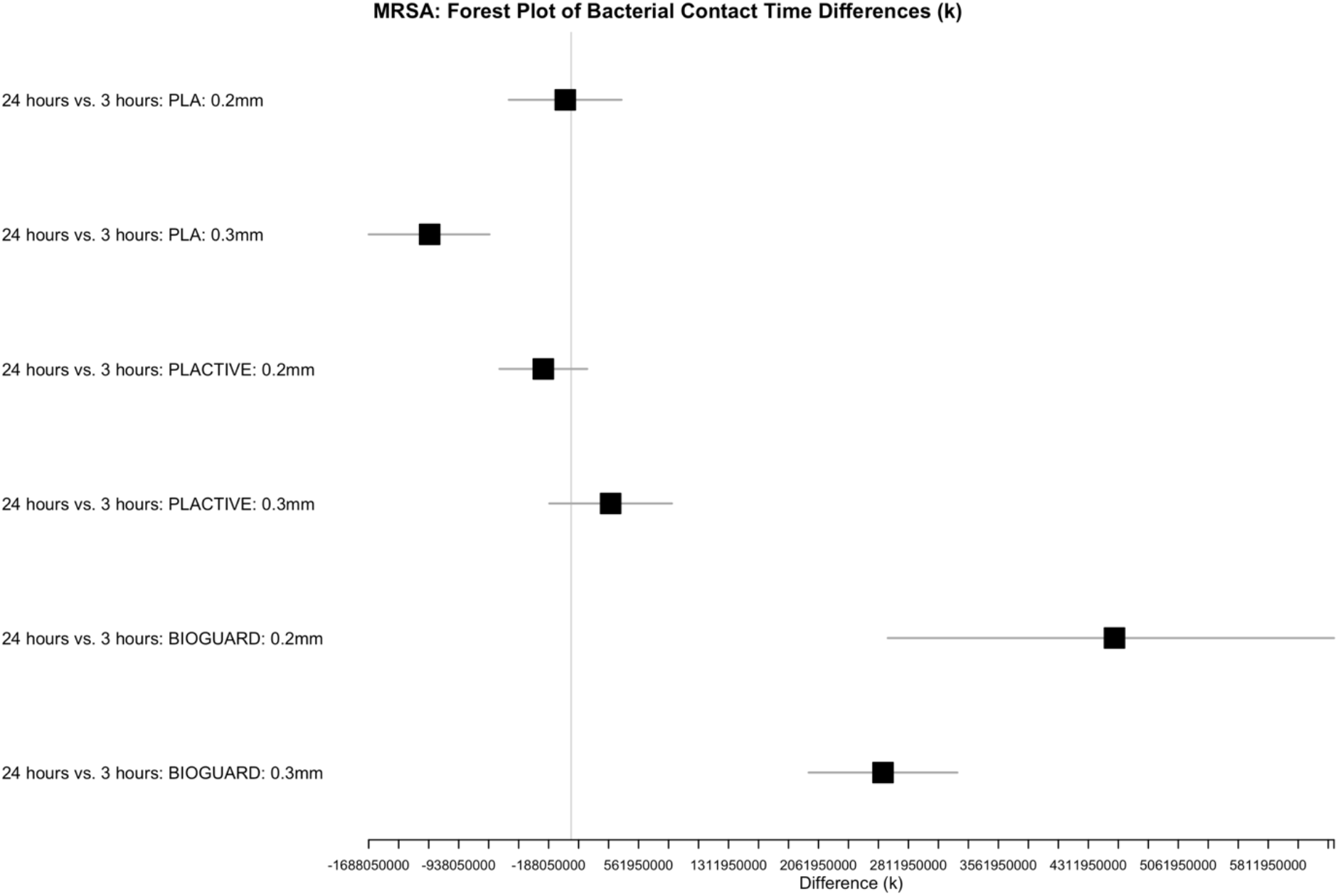
Forest plot illustrating the differences in k values between bacterial contact time for methicillin-resistant *Staphylococcus aureus*, categorized by material type and layer height. The plot displays the mean difference (k) along with scaled confidence intervals (Cis), where Cis exceeding a threshold value are scaled by a specified factor.

Both MRSA and *Staphylococcus aureus* exhibited increased carrying capacity when using BIOGUARD. Specifically, MRSA with BIOGUARD 0.3mm showed a significant increase in carrying capacity for the long bacterial contact time (2.6e9 [9.0e8, 4.3e9]), resulting in higher CFU/mL compared to shorter 3-hour contact time. Similarly, *Staphylococcus aureus* demonstrated a similar increase in carrying capacity with BIOGUARD 0.2mm (1.11e9 [4.50e8, 1.77e9]), resulting in higher CFU/mL during prolonged bacterial contact times compared to shorter 3-hour contact times. Interestingly, MRSA had a decreased carrying capacity for the long bacterial contact time for PLA at 0.3mm (-1.18e9 [-1.72e9, -6.49e8]) (Supplement Figures 8-14).

### Layer Height Impact on Growth Rate and Carrying Capacity

Our results indicated variability in the difference between the two layer heights, 0.2mm and 0.3mm, for carrying capacity, particularly at the shorter 3-hour bacterial contact time. When using a layer height of 0.2mm for PLA and BIOGUARD, *Klebsiella pneumonia* exhibited a higher CFU/mL (7.12e8 [1.19e8, 1.31e9]) and a lower CFU/mL (-1.17e9 [-2.22e9, -1.11e8]) respectively, indicating an increase and decrease in carrying capacity compared to the 0.3mm layer height. Regarding *Escherichia coli*, PLA (1.44e9 [8.45e8, 2.04e9]) and PLACTIVE (1.58e9 [8.40e8, 2.32e9]) showed an increase in carrying capacity when using 0.2mm layer height. Interestingly, both MRSA (-9.0e8 [-1.48e9, -3.17e8]) and *Staphylococcus aureus* (-5.74e8 [-9.97e8, -1.51e8]) had a decrease in carrying capacity when using PLA at 0.2mm layer height. (Figure 5, Supplement Figures 15-21)

**Figure 5:**
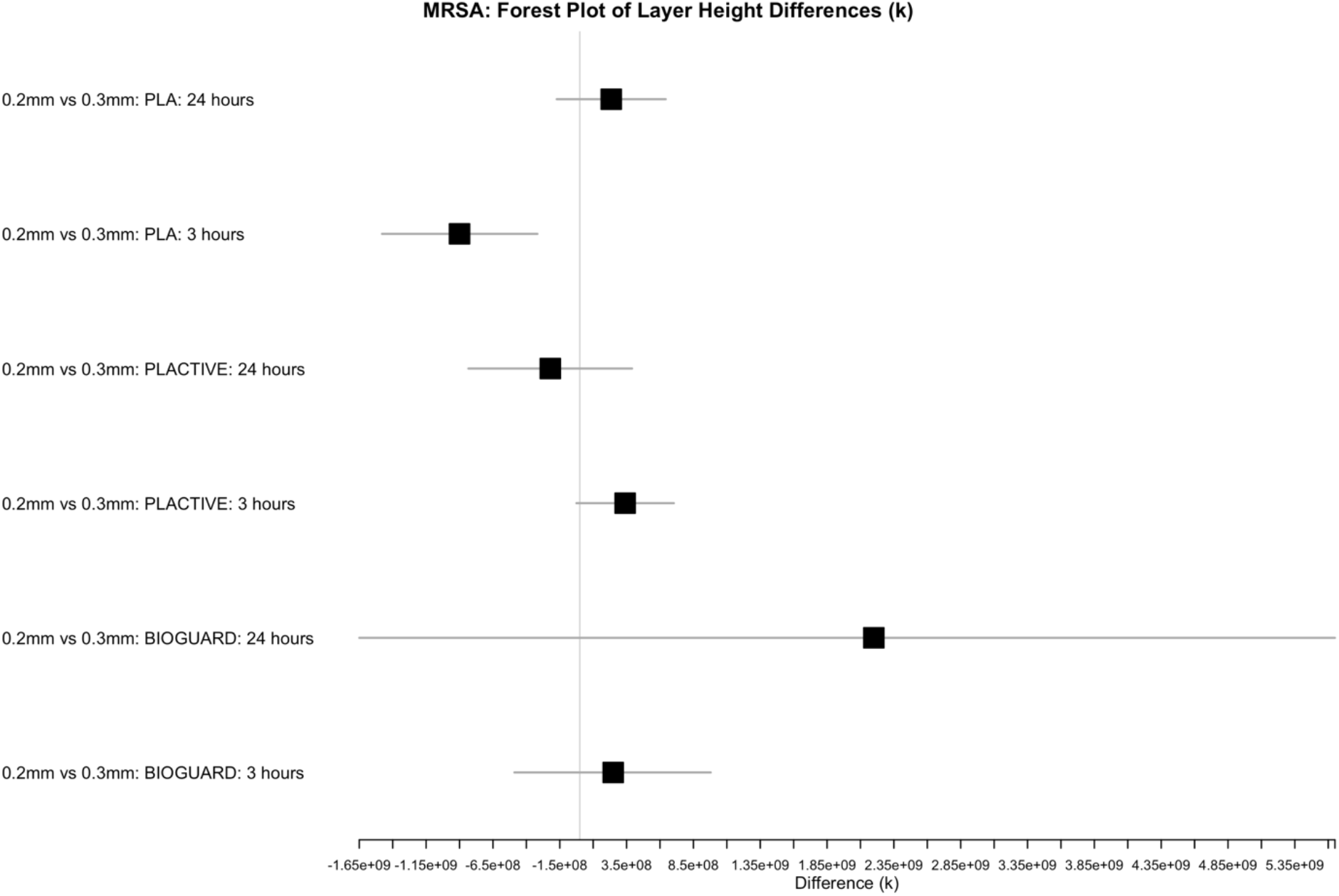
Forest plot illustrating the differences in k values between layer height for methicillin-resistant *Staphylococcus aureus*, categorized by material type and bacterial contact time. The plot displays the mean difference (k) along with scaled confidence intervals (Cis), where Cis exceeding a threshold value are scaled by a specified factor.

## DISCUSSION

In this study, we investigated the occurrence of bacterial contamination on polylactic acid (PLA) and evaluated the efficacy of disinfection measures, including an alternative solution-antimicrobial material. Our multifactorial analysis revealed significant effects of material type and bacterial species on colony-forming units (CFU), highlighting the complex dynamics of bacterial contamination in relation to different materials. The observed variations in CFU between PLA and alternative antimicrobial materials, such as PLACTIVE and BIOGUARD, underscore the importance of material selection in mitigating bacterial growth.

It is important to note that while PLACTIVE and BIOGUARD aim to address bacterial contamination through the addition of antimicrobial compounds into PLA, our results suggest the need for further optimization of their formulations to enhance antimicrobial efficacy. Copper3D, the makers of PLACTIVE, revealed that the antimicrobial properties were tested solely on the copper additive; it is unclear if the final product was included in the testing [26]. As our study showed, the efficacy of PLACTIVE and BIOGUARD in reducing bacterial contamination has shown varied results, emphasizing the need for comprehensive testing of antimicrobial additives in the final product. Moreover, our findings suggest that either additives such as copper and silver need to be more prominently present on the surface of the filament to ensure direct contact with potential bacteria, or there needs to be an increase in copper and silver ions in the filament. This highlights the importance of optimizing additive distribution within the filament matrix to enhance antimicrobial efficacy.

Filaments with much higher levels of metallic additives are available. For example, Prusa Research offers a tungsten filament to enable the fabrication of 3D-printed material with radiation-shielding properties. This material is 75% tungsten incorporated into a polyethylene terephthalate glycol (PETG) filament – another commonly used material. Such a high metallic content comes with considerable challenges; however, it requires a hardened steel nozzle vs. the brass or stainless-steel nozzles used on most 3D printers. It is also considerably more expensive than standard PETG, with a kilogram of filament costing $229 and providing ∼ 100m of filament, compared to PETG, costing approximately a tenth that amount for triple the length of the filament [27]. Increasing copper or silver ions in filaments may lead to the same challenges presented as tungsten and compromise the relative ease of printing of PLACTIVE and BIOGUARD. Companies that produce 3D printer filaments will need to incorporate further testing to elucidate potential challenges that come with increased metal in these filaments.

Multiple studies have shown the effectiveness of both copper and silver in reducing bacterial populations [19]. What’s more, copper’s antimicrobial efficacy rises in direct proportion to its concentration. The durability of stainless steel coated with a silver-zeolite matrix diminished over repeated washes, varying based on the application method of the matrix. Nevertheless, this approach effectively reduced bacteria by 87% after 4 hours and maintained a 100% reduction after 24 hours. Complete surface coverage with either metal or their alloys has resulted in a significant decrease in bacterial contamination [6], [7], [12], [18], [19], [28], [29]. Additionally, Vasiliev et al. identified a synergetic effect of mixed copper and silver nanoparticles due to their complementary mechanisms of action [30]. This further supports the importance of contact with copper and silver for efficiently eliminating or reducing bacteria.

Interestingly, while bacterial contact time and disinfectant contact times did not show significant effects on CFU in our study, further exploration using non-linear models unveiled nuanced differences in growth rate and carrying capacity between material types. For instance, *Klebsiella pneumonia* exhibited decreased carrying capacity on PLA compared to BIOGUARD, suggesting differential susceptibility to bacterial colonization on distinct materials.

Although our investigation into the impact of layer height revealed variability in carrying capacity, especially at shorter bacterial contact times, the overall findings suggest that layer height may not significantly impact bacterial contamination levels in the long run. The observed differences in CFU between different layer heights underscore the importance of considering printing parameters in 3D printing processes. However, further analysis suggests that other factors may have a more substantial influence on bacterial growth dynamics.

Our findings also revealed notable differences in bacterial growth dynamics between long (24 hours) and short (3 hours) bacterial contact times. The consistently higher CFU observed at longer contact times underscores the importance of timely disinfection measures to prevent microbial proliferation, particularly in healthcare settings.

Notably, the efficacy of disinfection measures, such as 70% EtOH, in reducing CFU on PLA underscores the importance of proper disinfection protocols in controlling bacterial contamination. However, it is essential to note that bacterial population levels returned hours after disinfectant application. An alternative solution to 70% EtOH is bleach, widely recognized as the gold standard in disinfection [31], [32], [33]. While bleach has the potential to disinfect PLA effectively, it is crucial to determine the appropriate concentration to ensure efficacy while also considering the possibility of material degradation. A balance must be struck between effective disinfection and preserving material integrity. Based on our findings, we recommend continuous disinfection with 70% EtOH or considering PLA as a single-use item to minimize the risk of bacterial contamination.

Our study provides valuable insights into the factors influencing bacterial contamination on PLA and highlights the importance of material selection and disinfection protocols in mitigating microbial proliferation. Further research is warranted to explore additional factors influencing bacterial growth dynamics and to develop effective strategies for controlling bacterial contamination in 3D-printed materials.

## Acknowledgments

The funders had no role in study design, data collection, and interpretation or the decision to submit the work for publication.

## Financial Support

KCJ was supported by a HIRe Fellowship from the U.S. Centers for Disease Control and Prevention (U01CK000673). ETL was supported by a grant from the National Institutes of Health (R35GM147013).

## Conflicts of Interest

None

Thank you to the staff and TAs at the Franceschi Microscopy and Imaging Center for their assistance and training.

## SUPPLEMENTAL FIGURES

**Supplement Figure 1.**
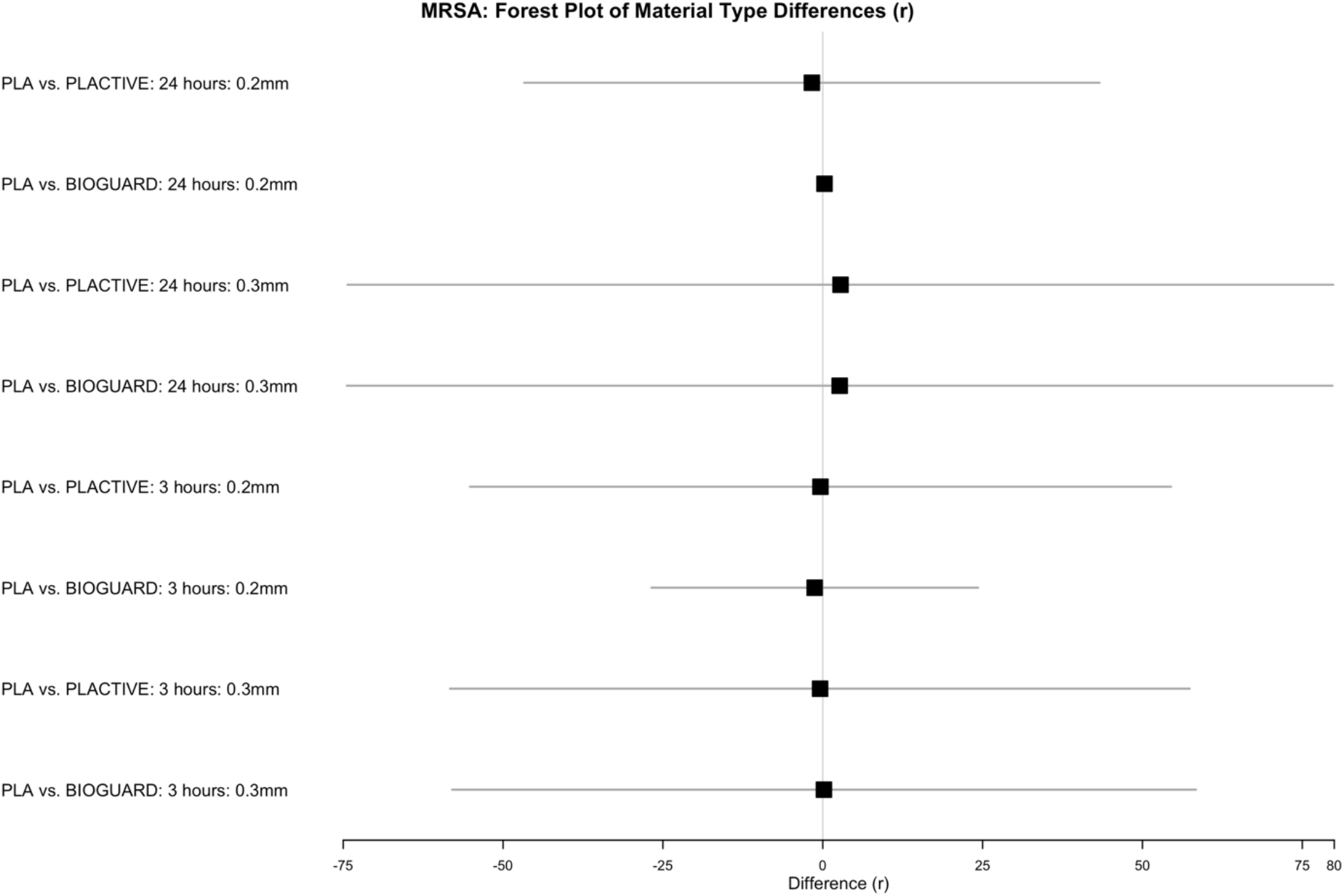
Forest plot illustrating the differences in r values between material types for methicillin-resistant *Staphylococcus aureus*, categorized by contact time and layer height. The plot displays the difference (r) along with scaled confidence intervals (CI), where CIs exceeding a threshold value are scaled by a specified factor.

**Supplement Figure 2.**
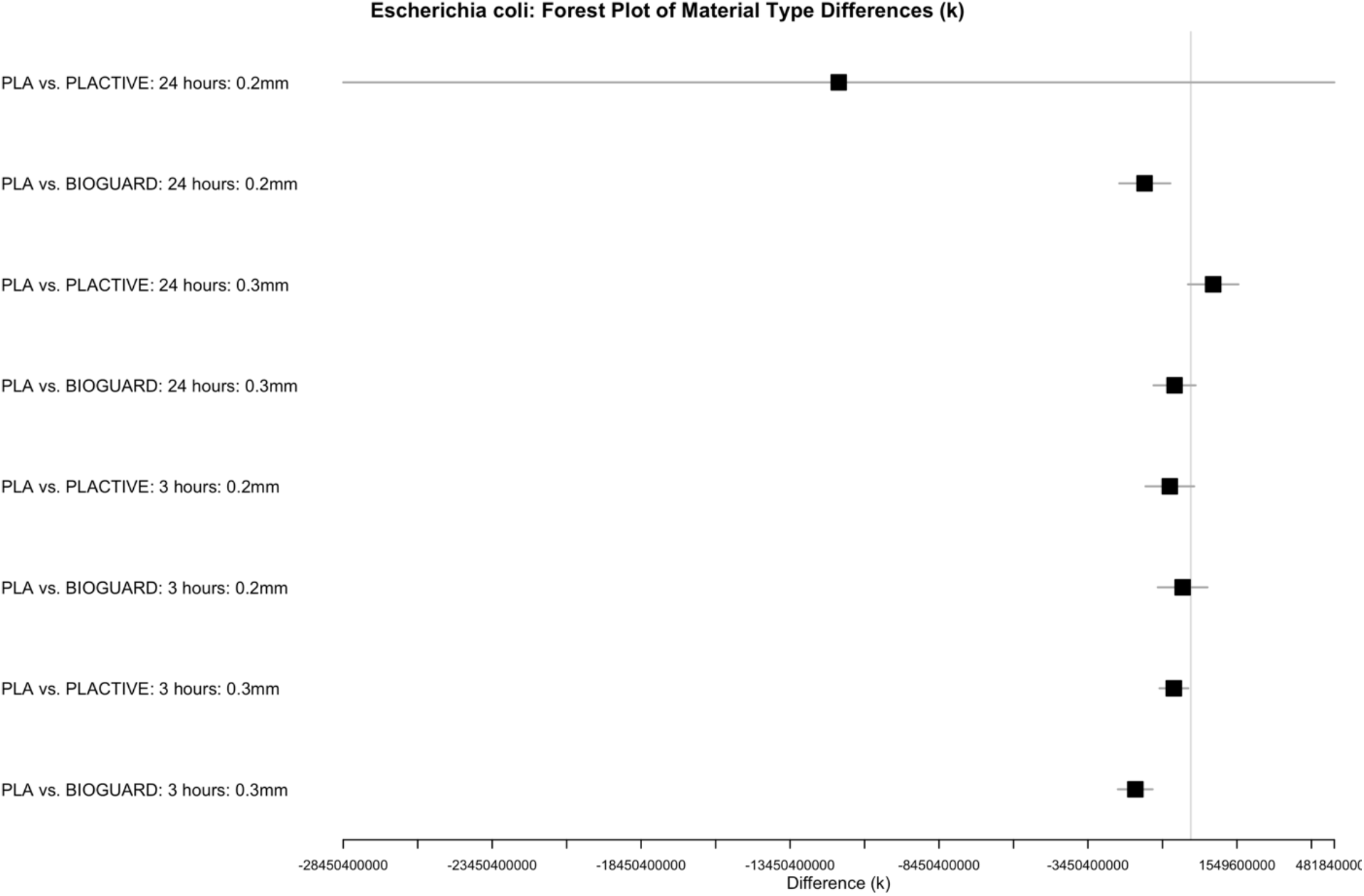
Forest plot illustrating the differences in k values between material types for *Escherichia coli*, categorized by contact time and layer height. The plot displays the difference (k) along with scaled confidence intervals (CI), where CIs exceeding a threshold value are scaled by a specified factor.

**Supplement Figure 3.**
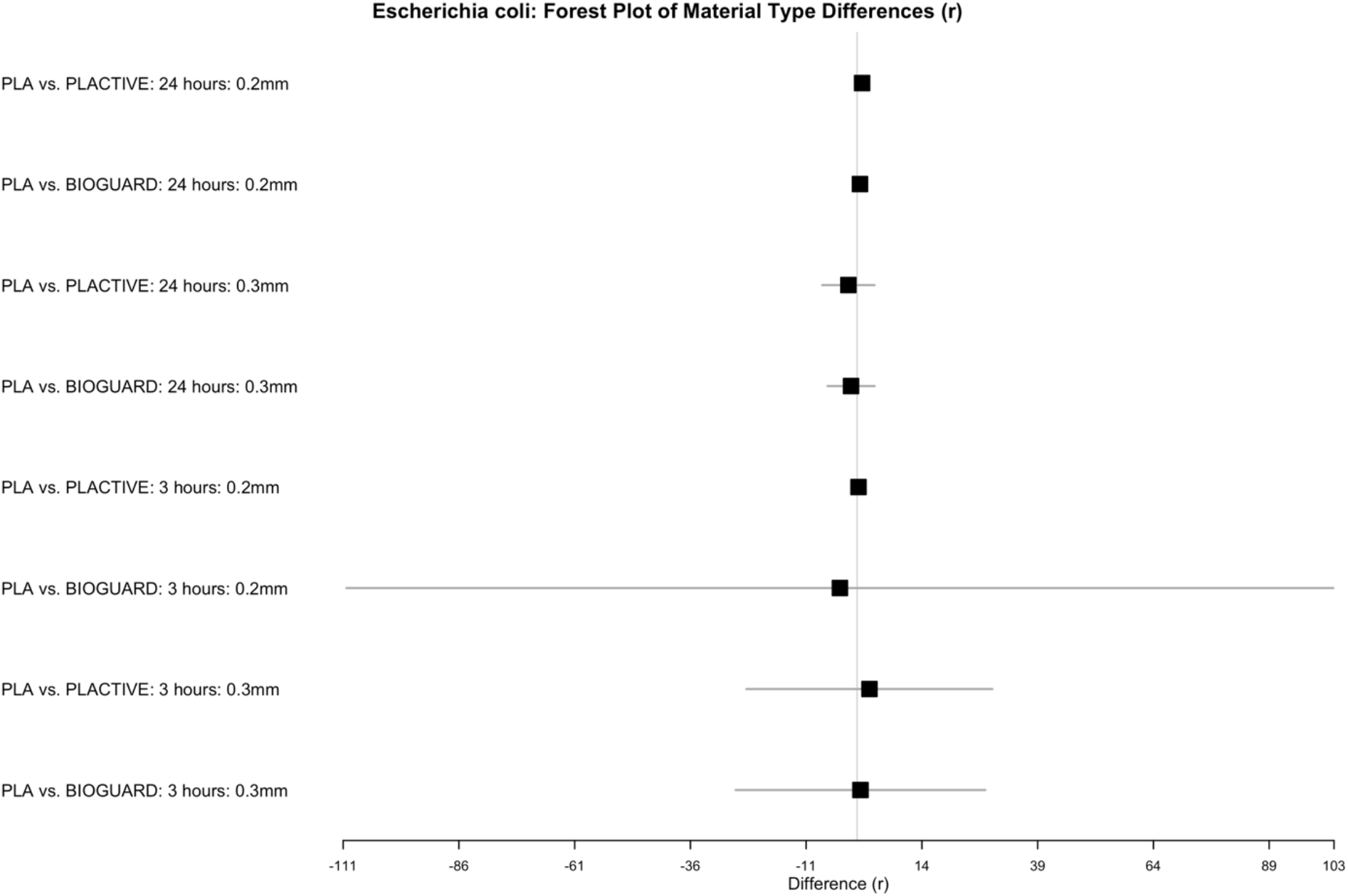
Forest plot illustrating the differences in r values between material types for *Escherichia coli*, categorized by contact time and layer height. The plot displays the difference (r) along with scaled confidence intervals (CI), where CIs exceeding a threshold value are scaled by a specified factor.

**Supplement Figure 4.**
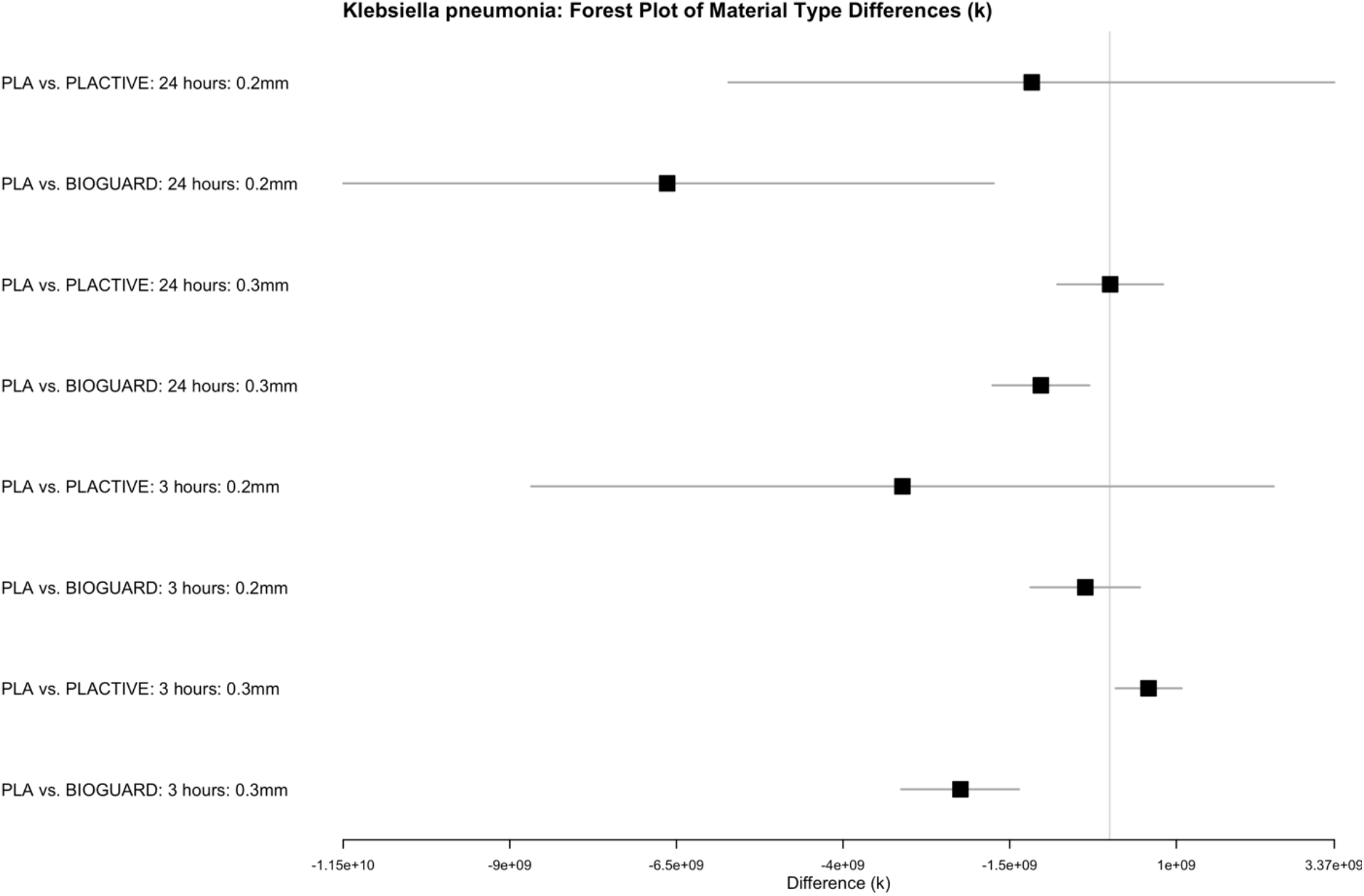
Forest plot illustrating the differences in k values between material types for *Klebsiella pneumonia*, categorized by contact time and layer height. The plot displays the difference (k) along with scaled confidence intervals (CI), where CIs exceeding a threshold value are scaled by a specified factor.

**Supplement Figure 5.**
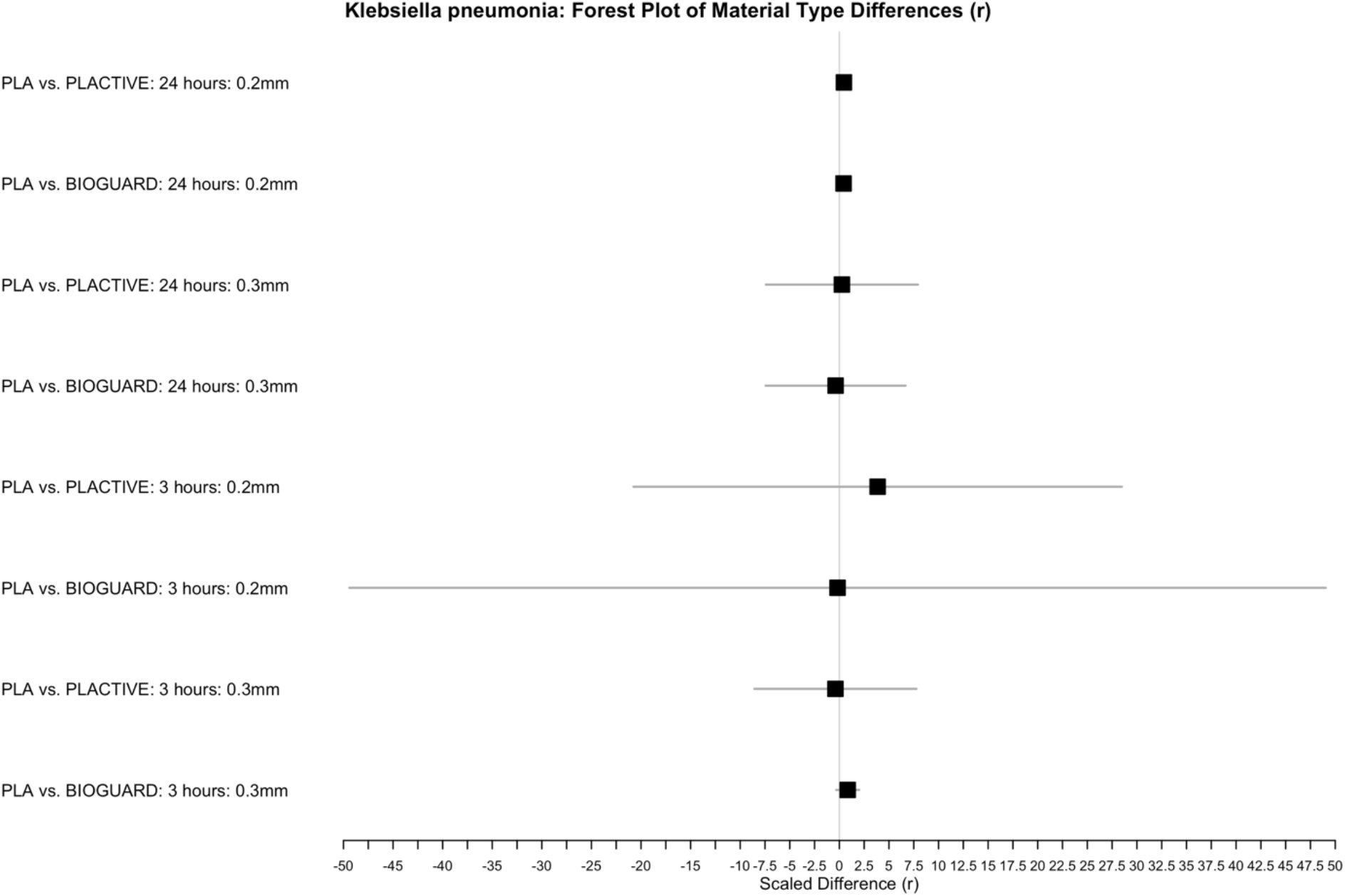
Forest plot illustrating the differences in r values between material types for *Klebsiella pneumonia*, categorized by contact time and layer height. The plot displays the difference (r) along with scaled confidence intervals (CI), where CIs exceeding a threshold value are scaled by a specified factor.

**Supplement Figure 6.**
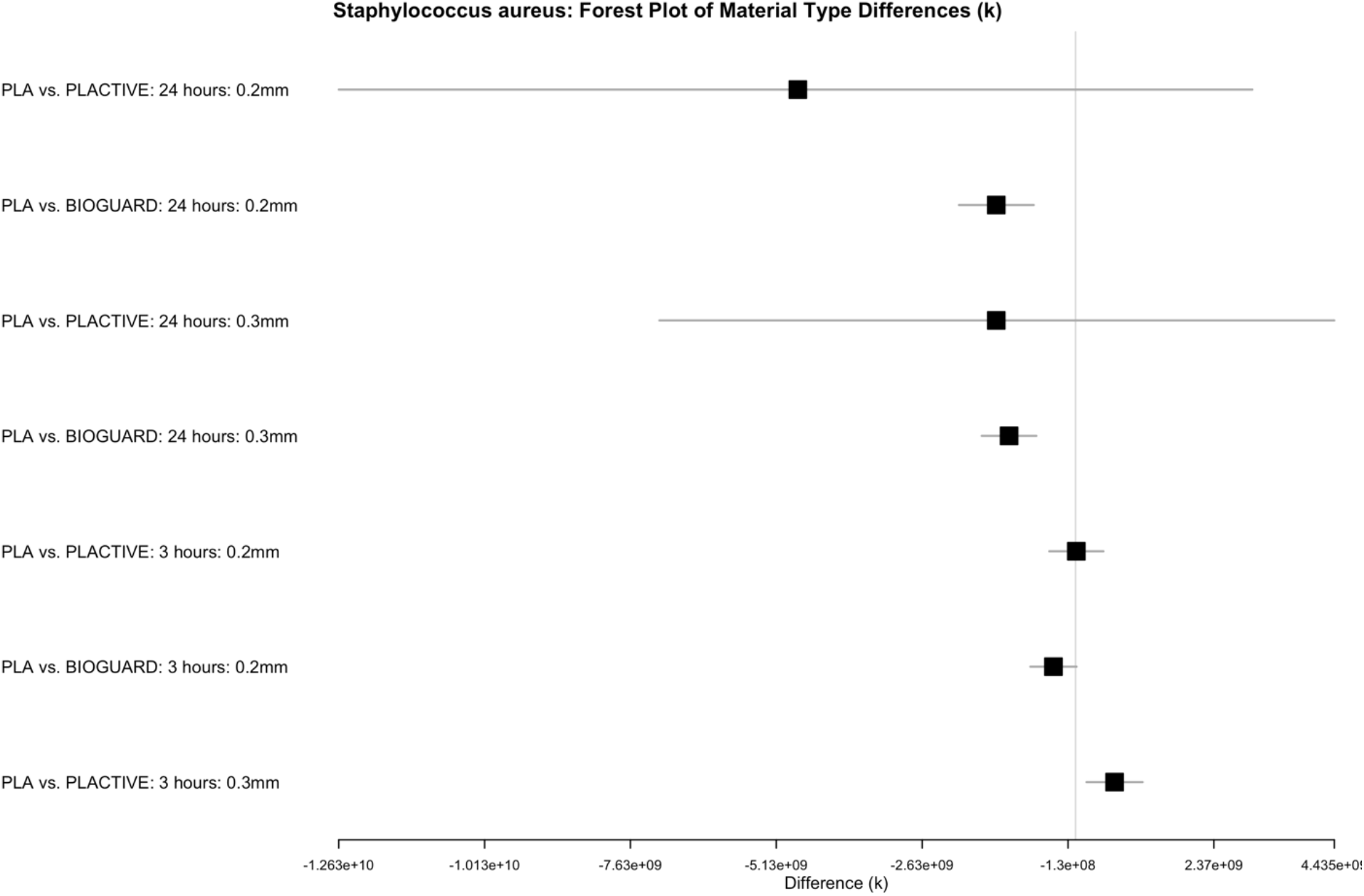
Forest plot illustrating the differences in k values between material types for *Staphylococcus aureus*, categorized by contact time and layer height. The plot displays the difference (k) along with scaled confidence intervals (CI), where CIs exceeding a threshold value are scaled by a specified factor. Values exceeding the threshold are omitted from the plot for clarity.

**Supplement Figure 7.**
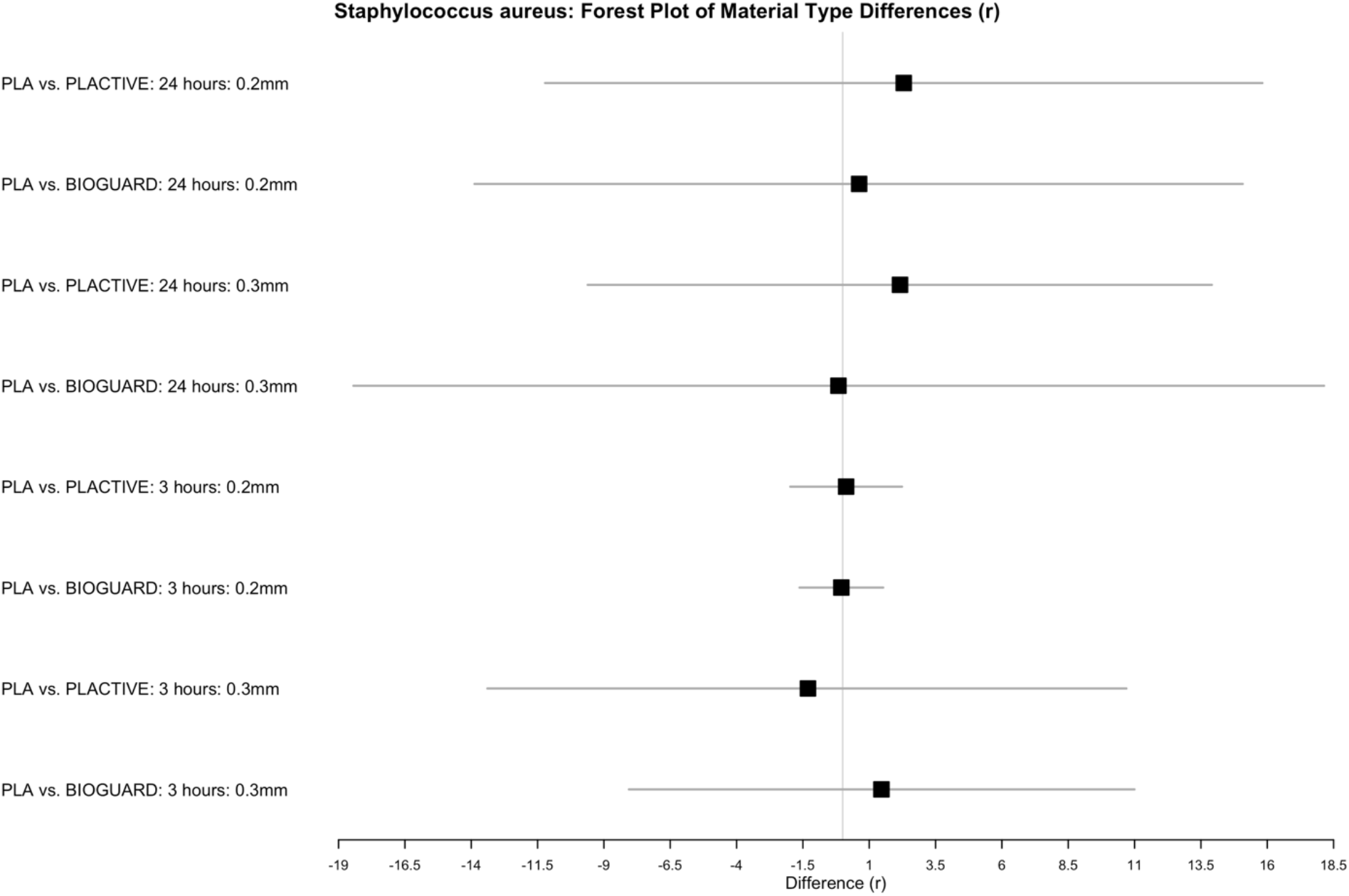
Forest plot illustrating the differences in r values between material types for *Staphylococcus aureus*, categorized by contact time and layer height. The plot displays the difference (r) along with scaled confidence intervals (CI), where CIs exceeding a threshold value are scaled by a specified factor.

**Supplement Figure 8.**
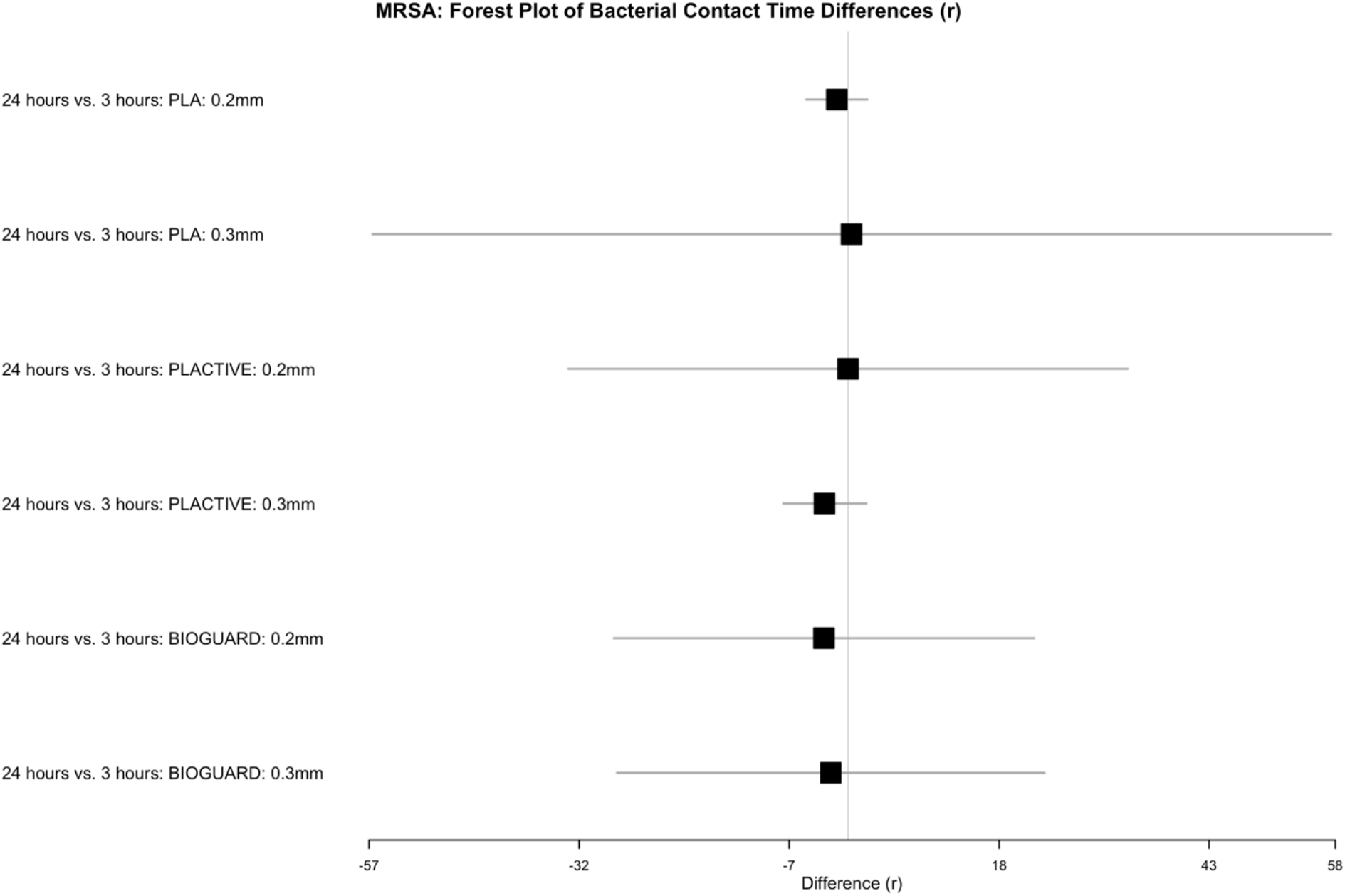
Forest plot illustrating the differences in r values between bacterial contact time for methicillin-resistant *Staphylococcus aureus*, categorized by material type and layer height. The plot displays the difference (r) along with scaled confidence intervals (CI), where CIs exceeding a threshold value are scaled by a specified factor.

**Supplement Figure 9.**
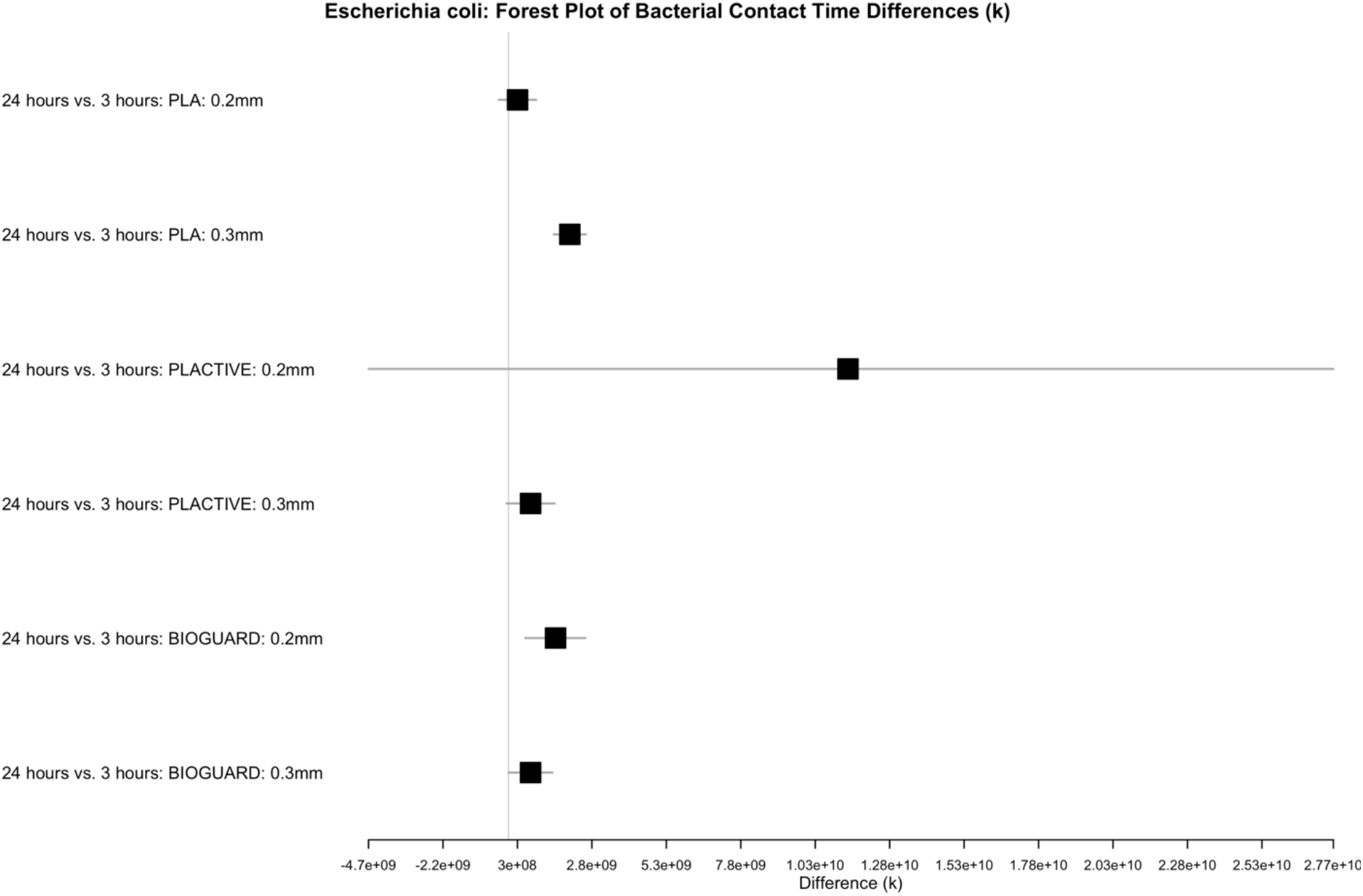
Forest plot illustrating the differences in k values between bacterial contact for *Escherichia coli*, categorized by material type and layer height. The plot displays the difference (k) along with scaled confidence intervals (CI), where CIs exceeding a threshold value are scaled by a specified factor.

**Supplement Figure 10.**
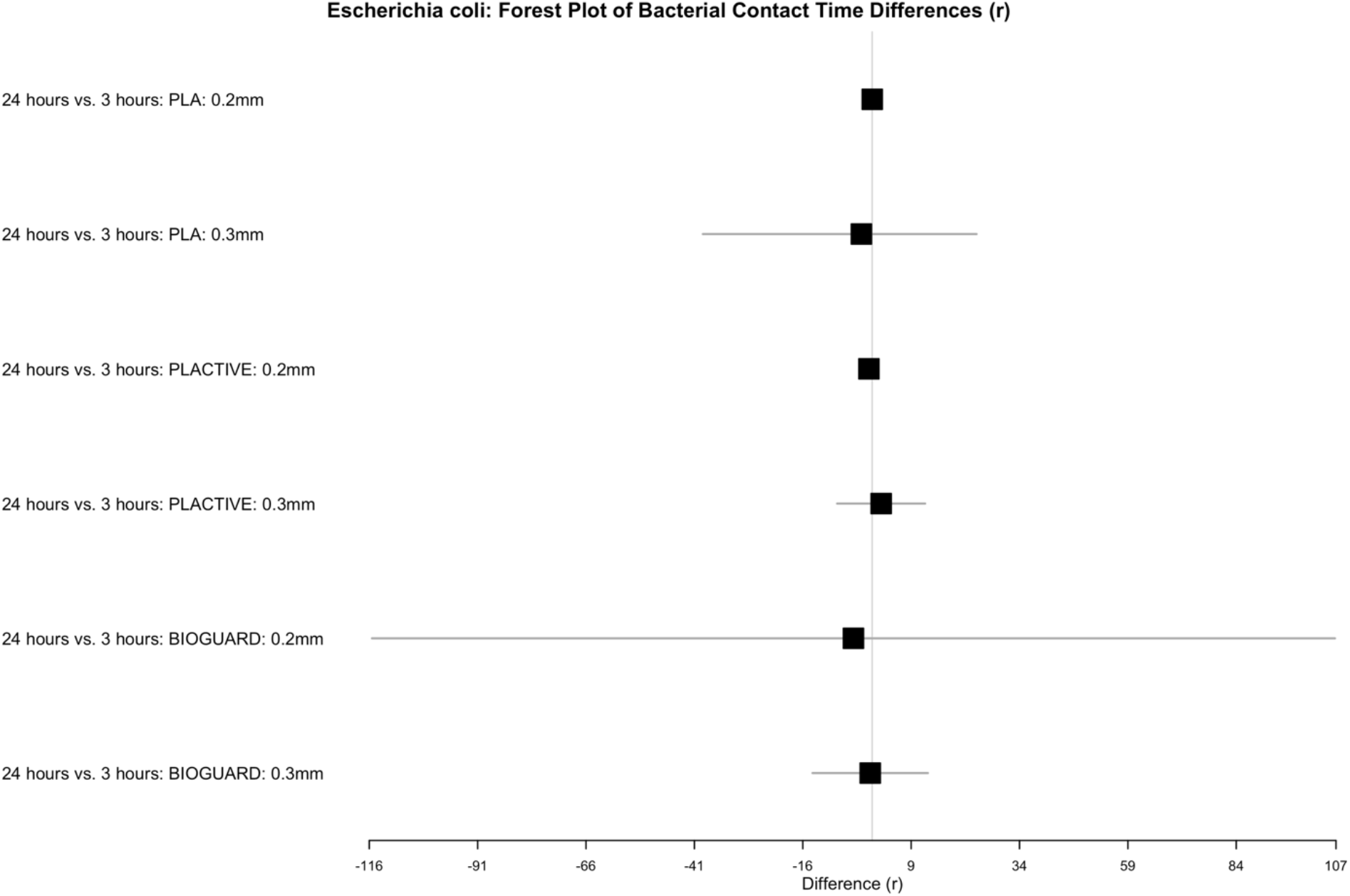
Forest plot illustrating the differences in r values between bacterial contact for *Escherichia coli*, categorized by material type and layer height. The plot displays the difference (r) along with scaled confidence intervals (CI), where CIs exceeding a threshold value are scaled by a specified factor.

**Supplement Figure 11.**
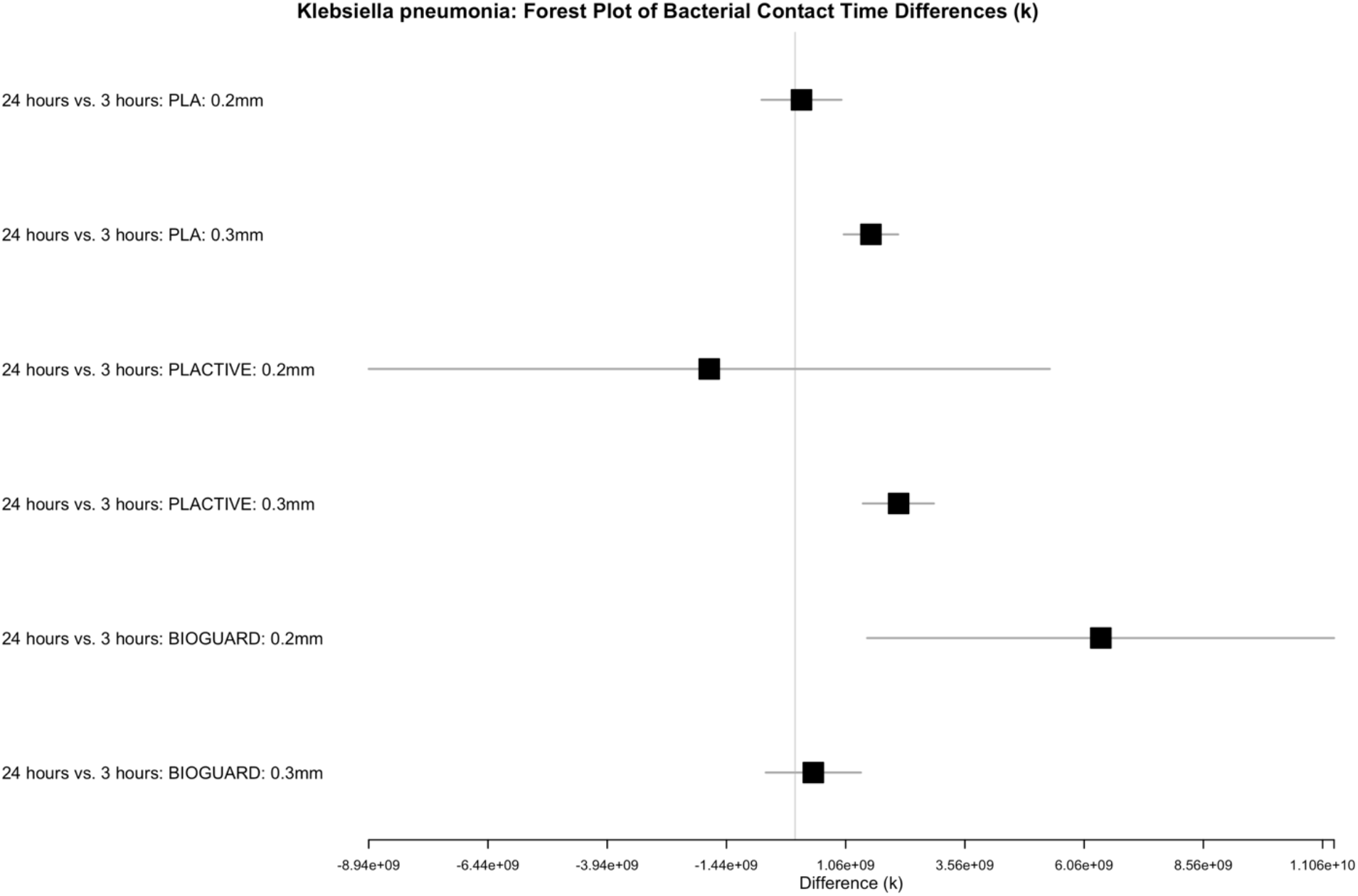
Forest plot illustrating the differences in k values between bacterial contact for *Klebsiella pneumonia*, categorized by material type and layer height. The plot displays the difference (k) along with scaled confidence intervals (CI), where CIs exceeding a threshold value are scaled by a specified factor.

**Supplement Figure 12.**
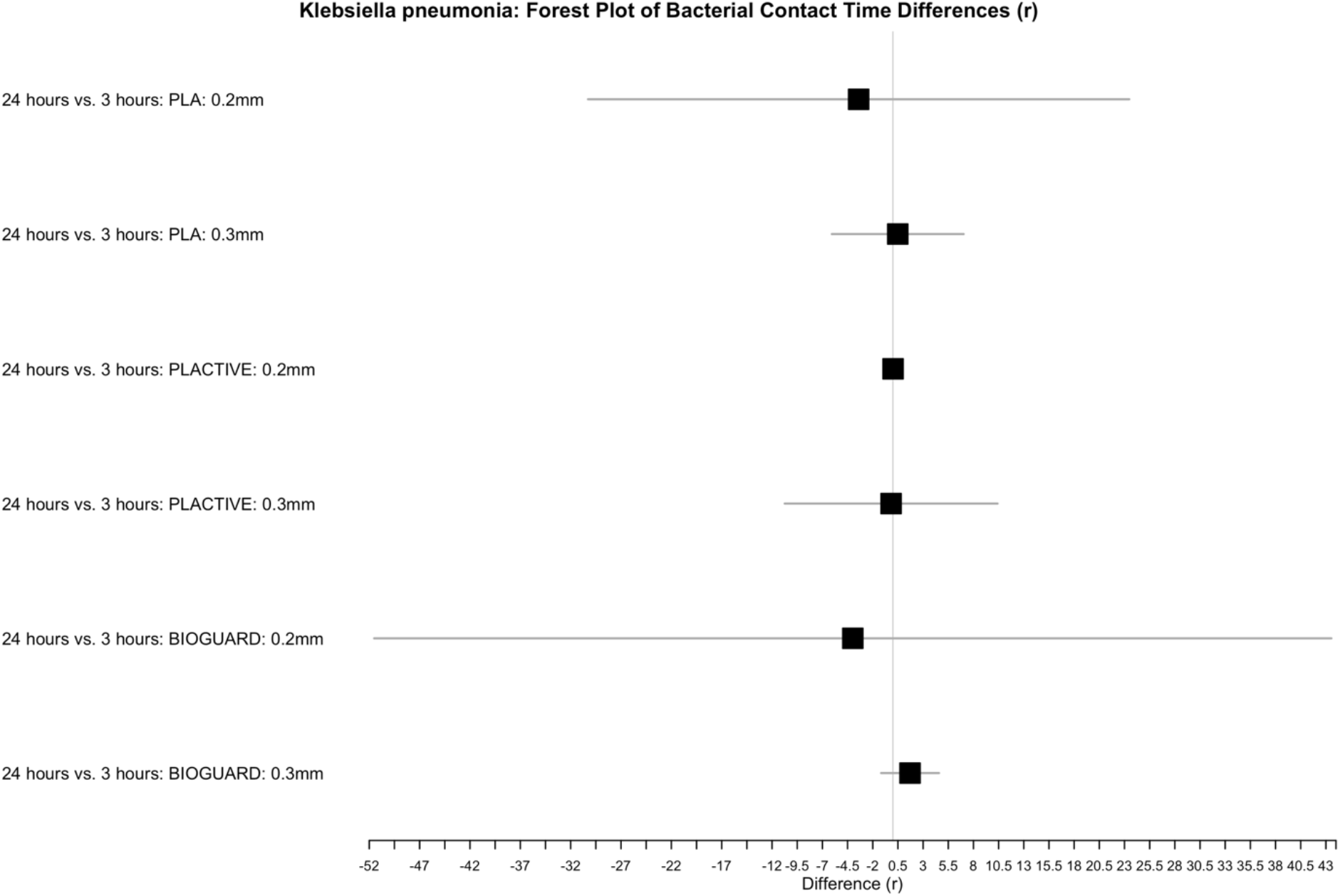
Forest plot illustrating the differences in r values between bacterial contact for *Klebsiella pneumonia*, categorized by material type and layer height. The plot displays the difference (r) along with scaled confidence intervals (CI), where CIs exceeding a threshold value are scaled by a specified factor.

**Supplement Figure 13.**
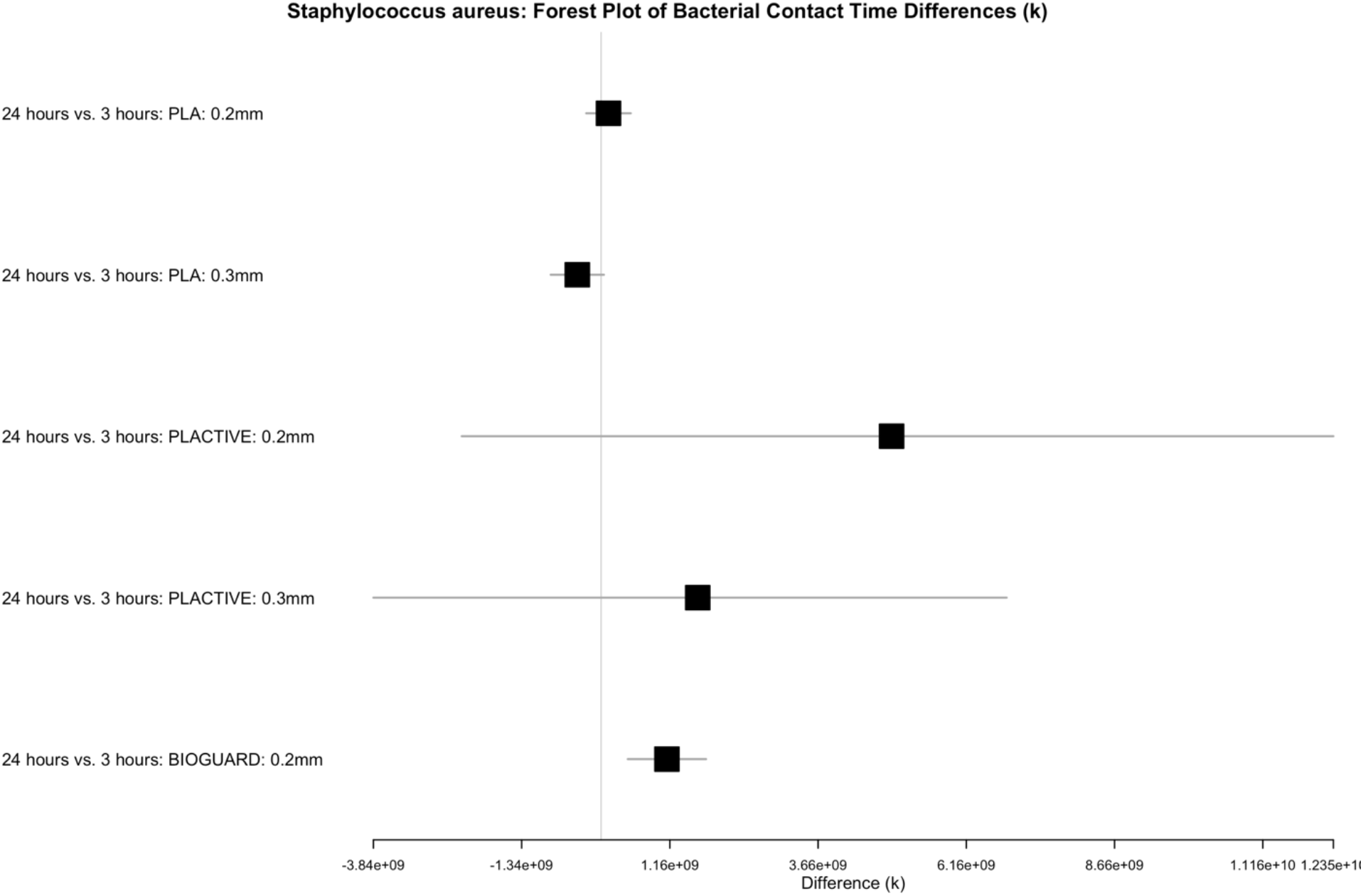
Forest plot illustrating the differences in k values between bacterial contact for *Staphylococcus aureus*, categorized by material type and layer height. The plot displays the difference (k) along with scaled confidence intervals (CI), where CIs exceeding a threshold value are scaled by a specified factor.

**Supplement Figure 14.**
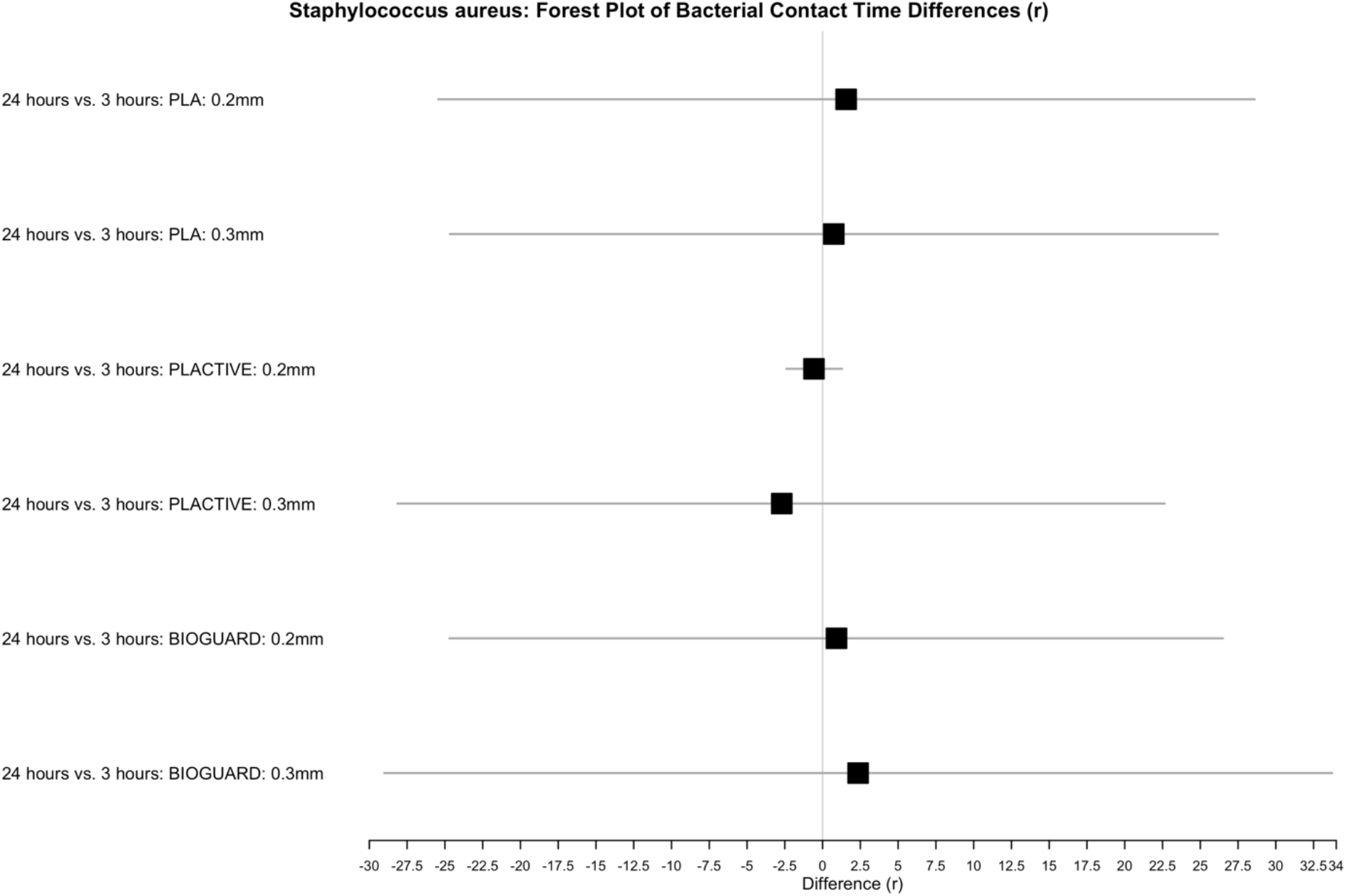
Forest plot illustrating the differences in r values between bacterial contact for *Staphylococcus aureus,* categorized by material type and layer height. The plot displays the difference (r) along with scaled confidence intervals (CI), where CIs exceeding a threshold value are scaled by a specified factor.

**Supplement Figure 15.**
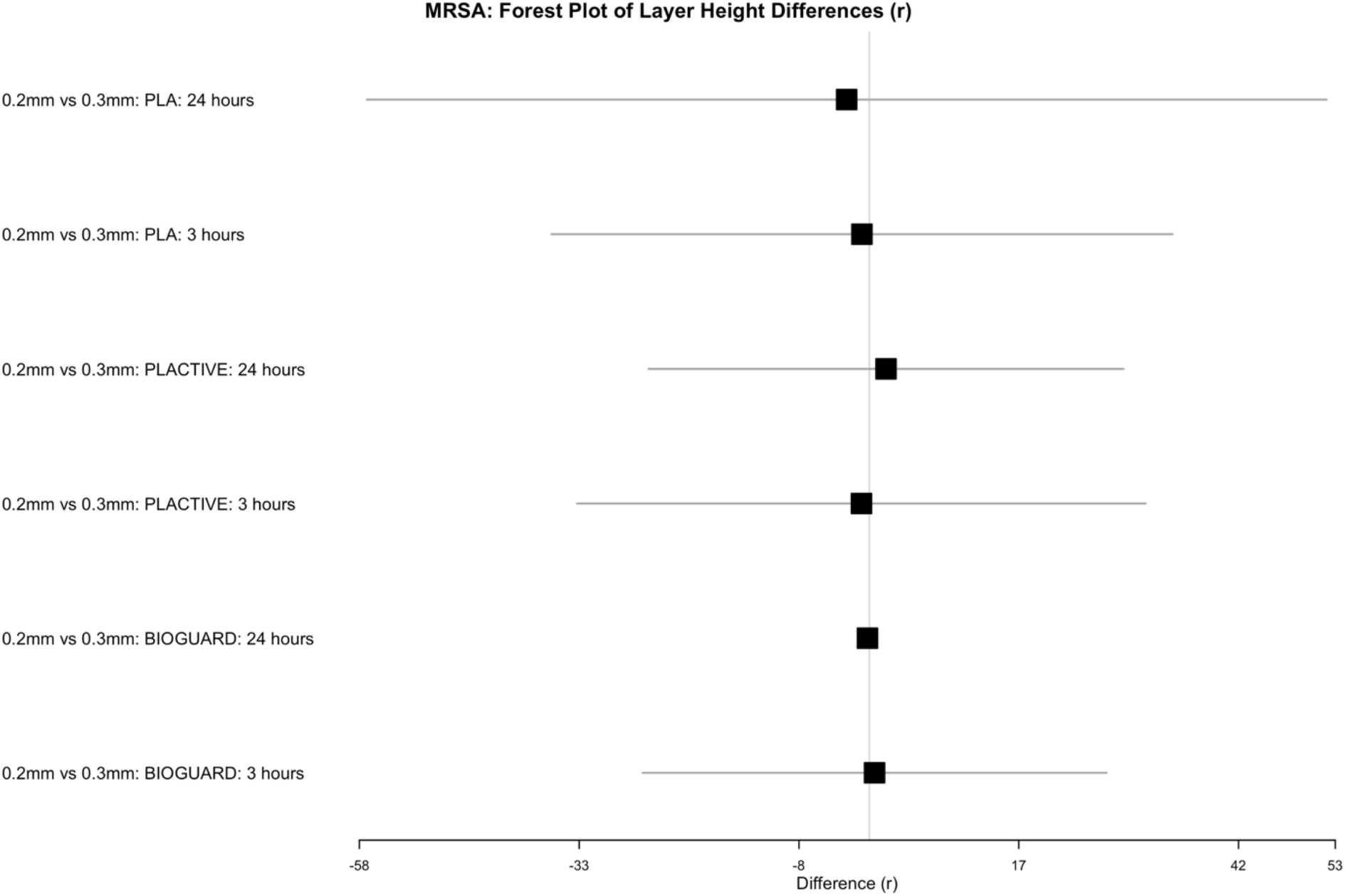
Forest plot illustrating the differences in r values between layer height for methicillin-resistant *Staphylococcus aureus*, categorized by material type and bacterial contact time. The plot displays the difference (r) along with scaled confidence intervals (CI), where CIs exceeding a threshold value are scaled by a specified factor.

**Supplement Figure 16.**
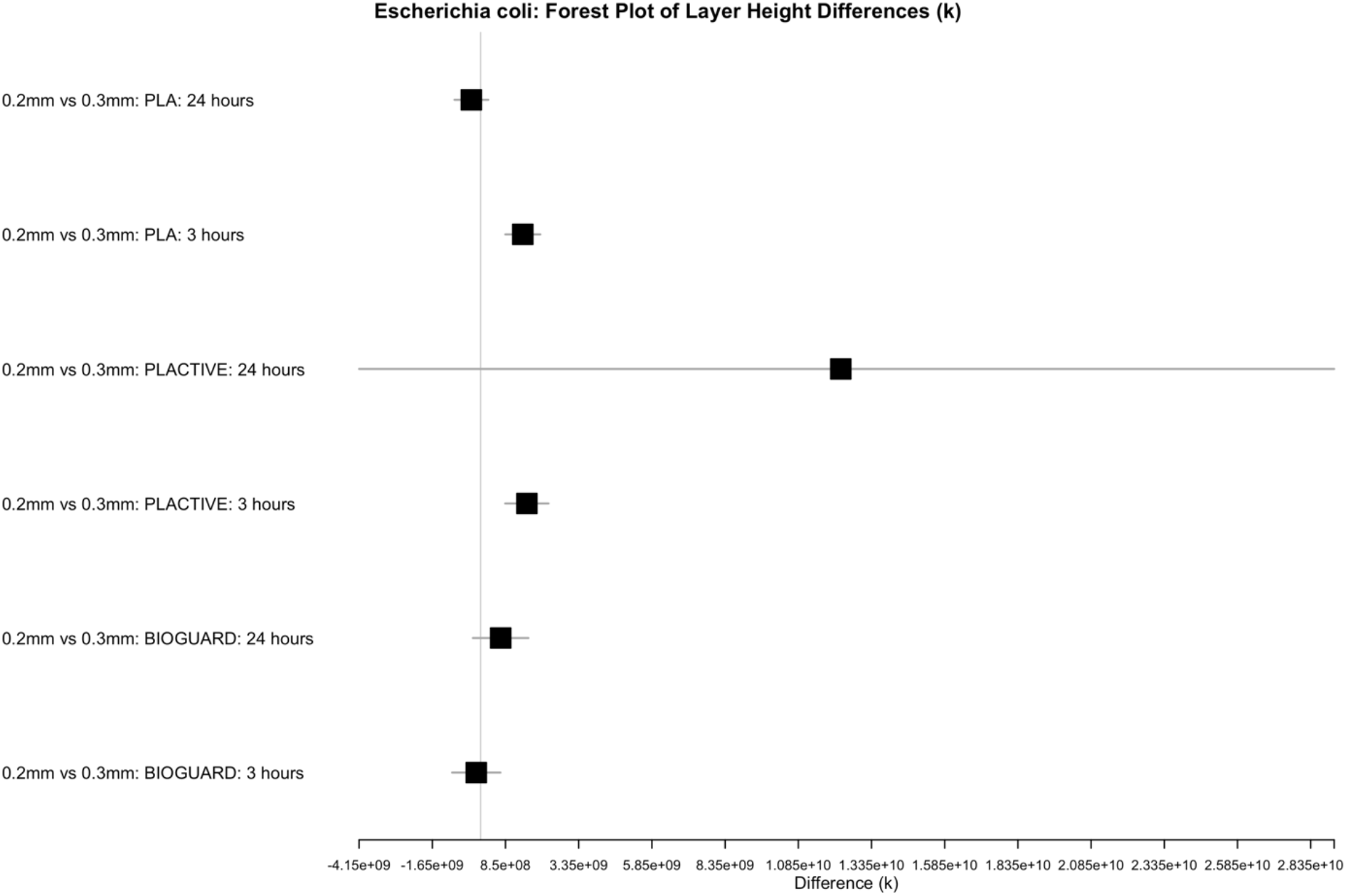
Forest plot illustrating the differences in k values between layer height for *Escherichia coli*, categorized by material type and bacterial contact time. The plot displays the difference (k) along with scaled confidence intervals (CI), where CIs exceeding a threshold value are scaled by a specified factor.

**Supplement Figure 17.**
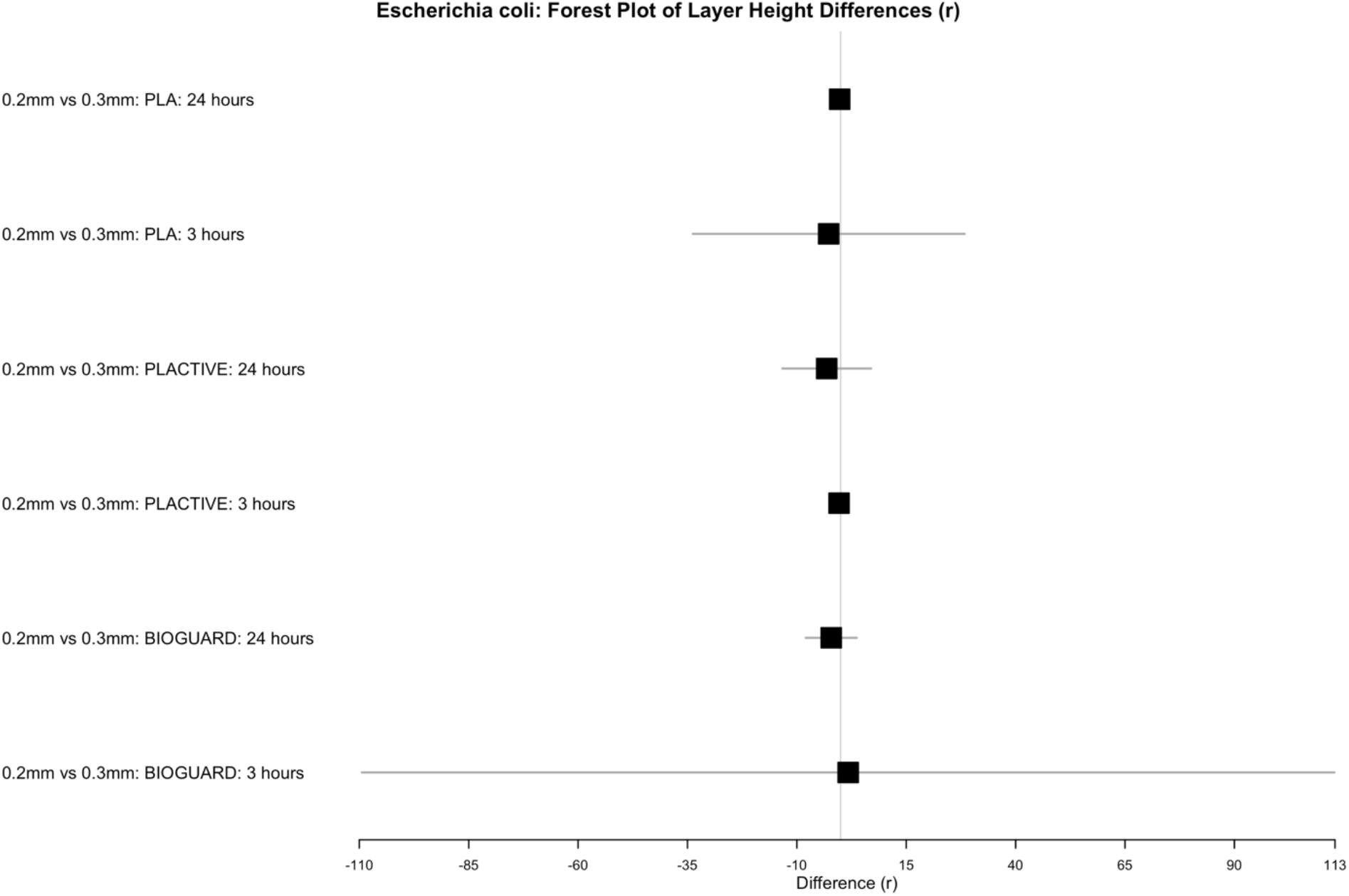
Forest plot illustrating the differences in r values between layer height for *Escherichia coli*, categorized by material type and bacterial contact time. The plot displays the difference (r) along with scaled confidence intervals (CI), where CIs exceeding a threshold value are scaled by a specified factor.

**Supplement Figure 18.**
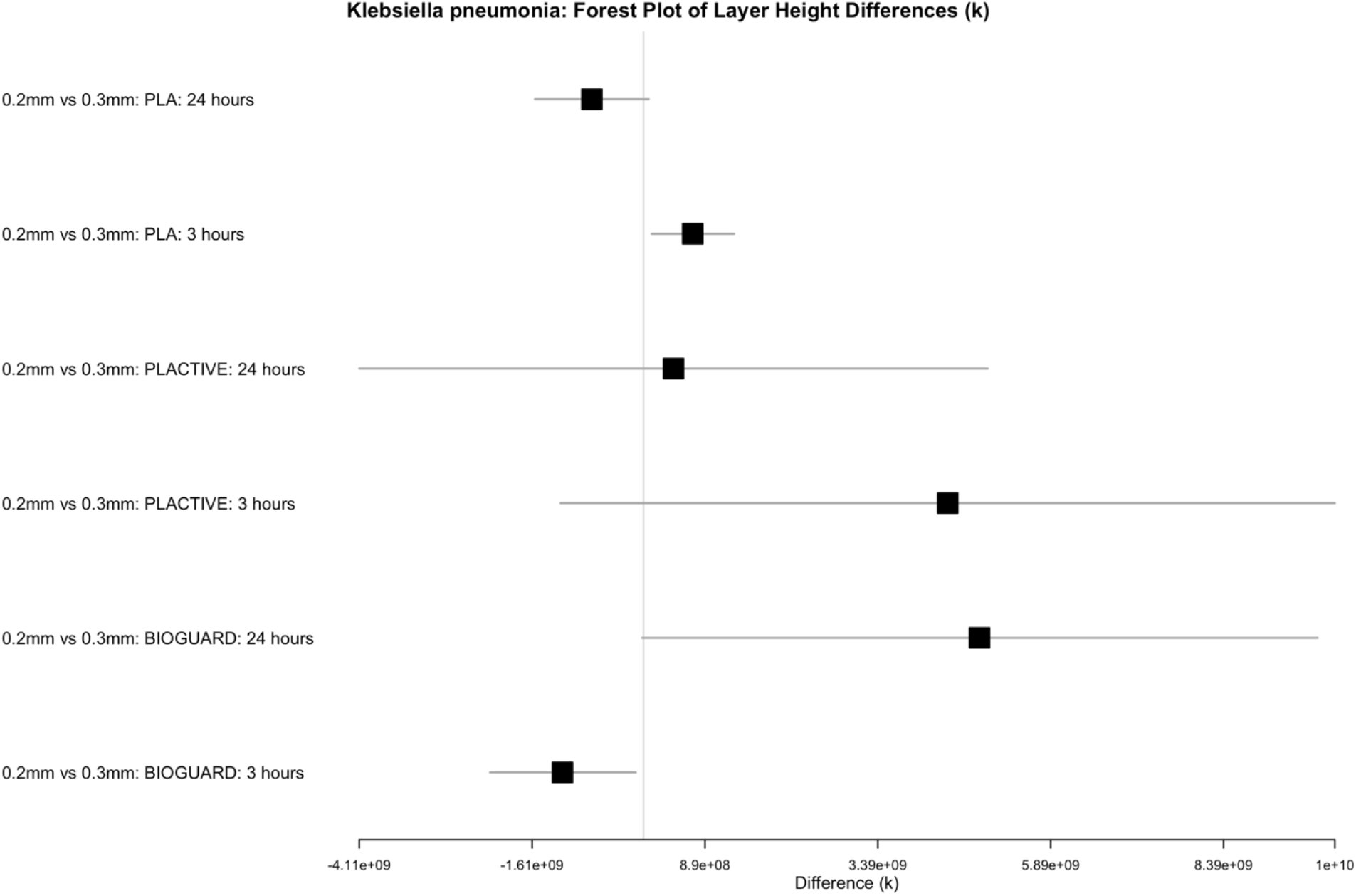
Forest plot illustrating the differences in k values between layer height for *Klebsiella pneumonia*, categorized by material type and bacterial contact time. The plot displays the difference (k) along with scaled confidence intervals (CI), where CIs exceeding a threshold value are scaled by a specified factor.

**Supplement Figure 19.**
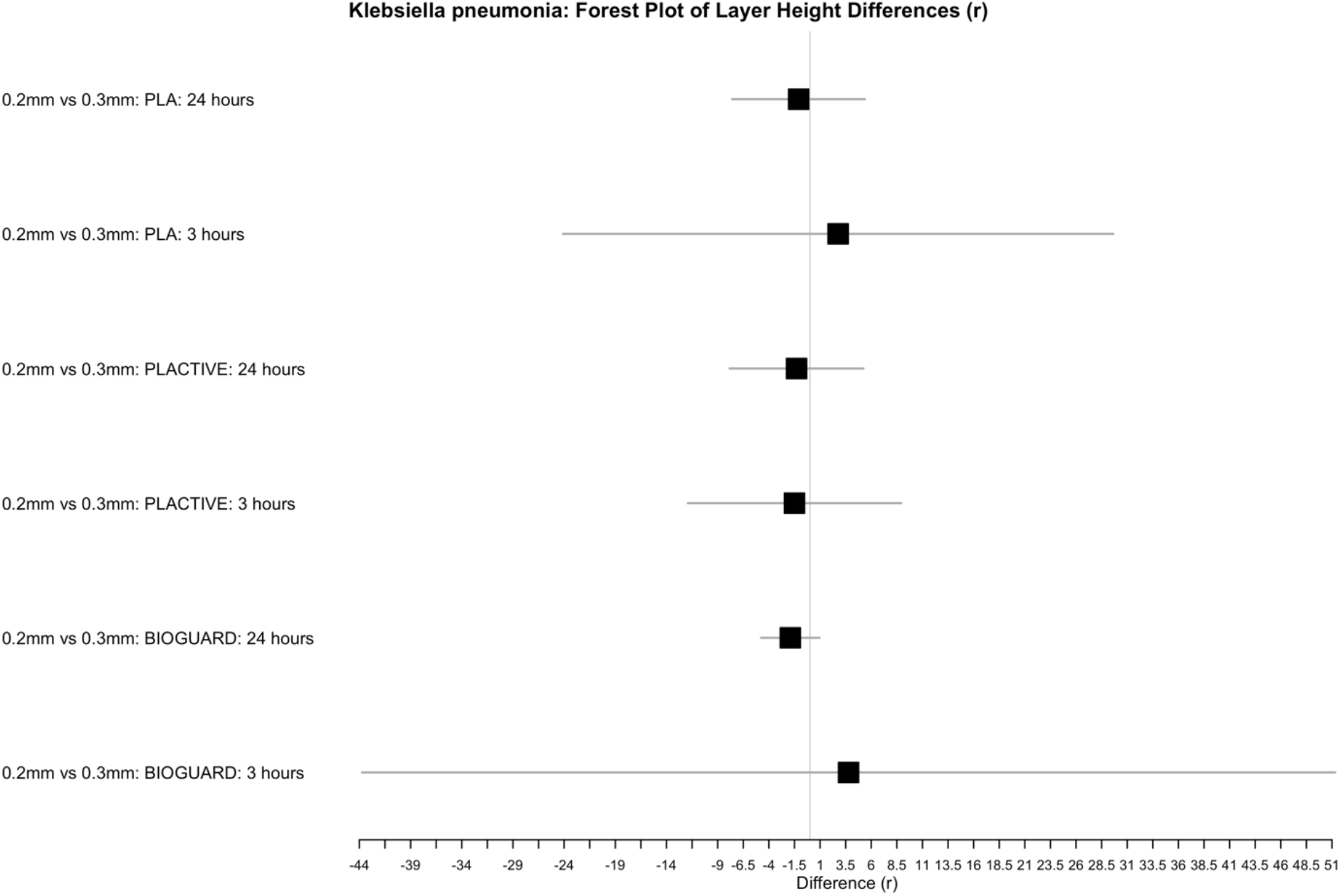
Forest plot illustrating the differences in r values between layer height for *Klebsiella pneumonia*, categorized by material type and bacterial contact time. The plot displays the difference (r) along with scaled confidence intervals (CI), where CIs exceeding a threshold value are scaled by a specified factor.

**Supplement Figure 20.**
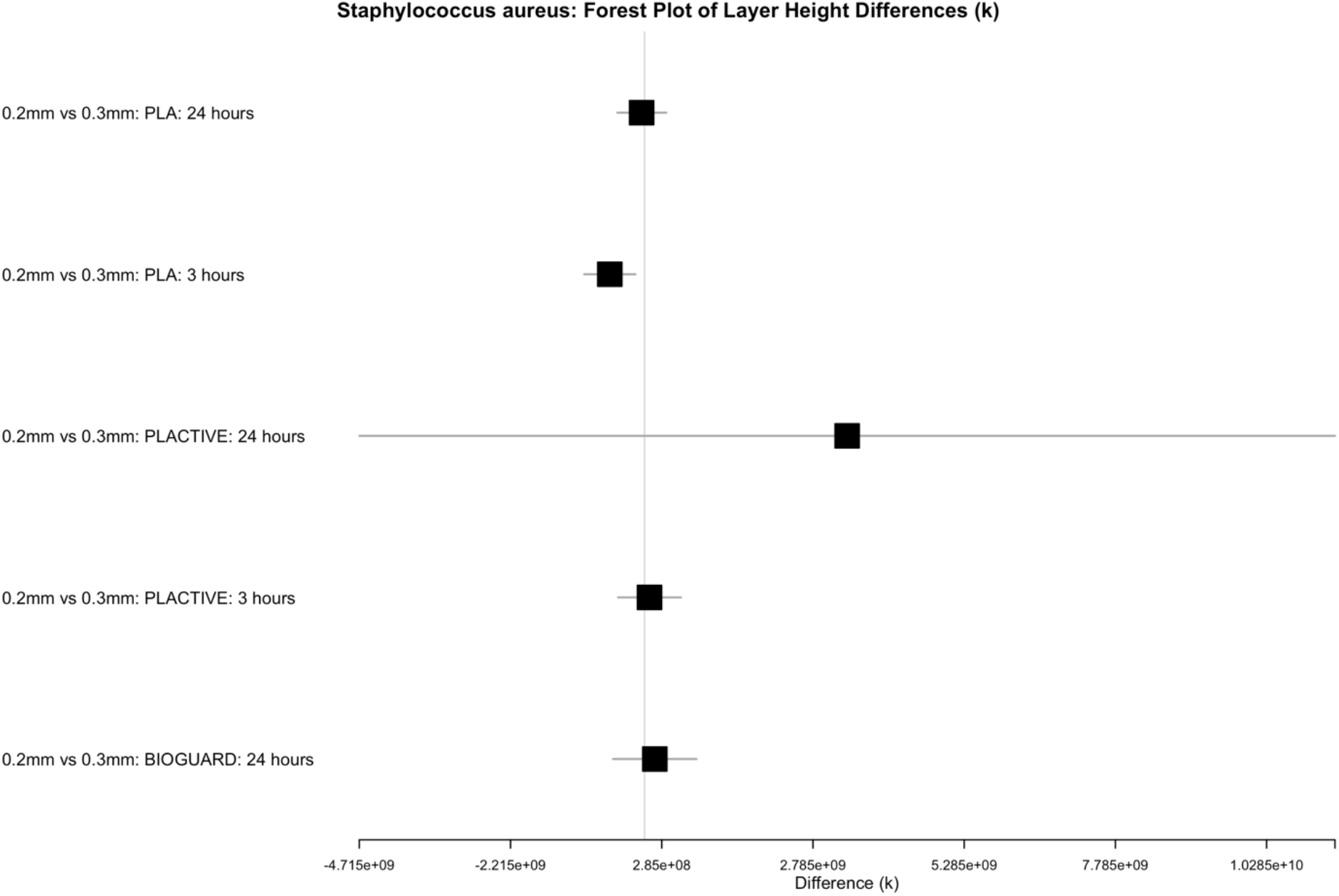
Forest plot illustrating the differences in k values between layer height for *Staphylococcus aureus*, categorized by material type and bacterial contact time. The plot displays the difference (k) along with scaled confidence intervals (CI), where CIs exceeding a threshold value are scaled by a specified factor. Values exceeding the threshold are omitted from the plot for clarity.

**Supplement Figure 21.**
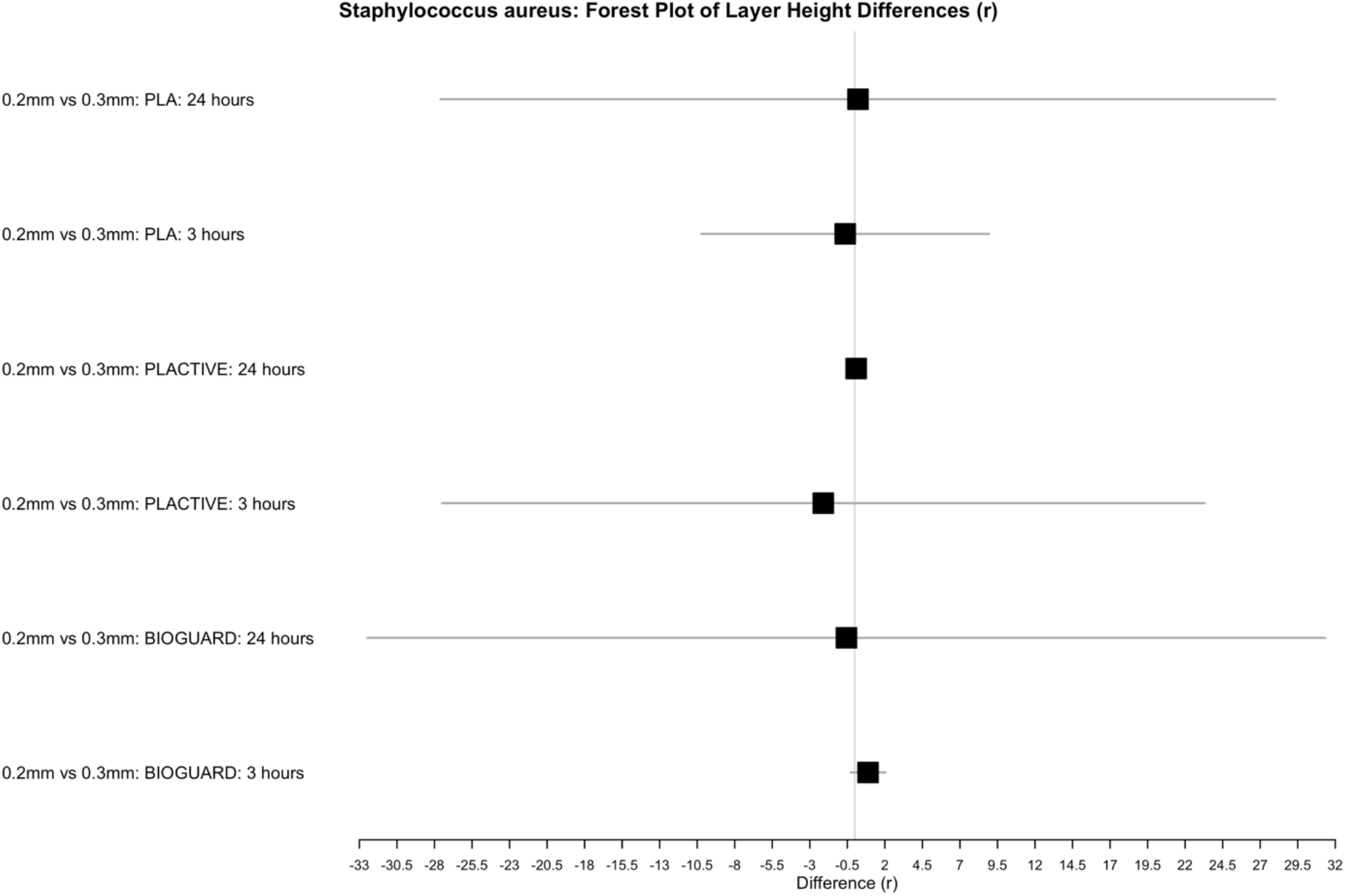
Forest plot illustrating the differences in r values between layer height for *Staphylococcus aureus*, categorized by material type and bacterial contact time. The plot displays the difference (r) along with scaled confidence intervals (CI), where CIs exceeding a threshold value are scaled by a specified factor.

## Notes

### Competing Interest Statement

The authors have declared no competing interest.

